# Peptide-enabled ribonucleoprotein delivery for CRISPR engineering (PERC) in primary human immune cells and hematopoietic stem cells

**DOI:** 10.1101/2024.07.14.603391

**Authors:** Srishti U Sahu, Madalena Castro, Joseph J Muldoon, Kunica Asija, Stacia K Wyman, Netravathi Krishnappa, Lorena de Oñate, Justin Eyquem, David N Nguyen, Ross C Wilson

**Affiliations:** Innovative Genomics Institute, University of California Berkeley, Berkeley, CA, USA; Department of Molecular and Cell Biology, University of California Berkeley, Berkeley, CA, USA; California Institute for Quantitative Biosciences at University of California Berkeley, Berkeley, CA, USA; Gladstone-UCSF Institute of Genomic Immunology, San Francisco, USA; Department of Medicine, University of California San Francisco, San Francisco, USA; Parker Institute for Cancer Immunotherapy, University of California San Francisco, San Francisco, CA, USA; Department of Microbiology and Immunology, University of California San Francisco, San Francisco, USA; UCSF Helen Diller Family Comprehensive Cancer Center, University of California San Francisco, USA; Institute for Human Genetics, University of California San Francisco, San Francisco, USA

## Abstract

Peptide-enabled ribonucleoprotein delivery for CRISPR engineering (PERC) is a new approach for *ex vivo* genome editing of primary human cells. PERC uses a single amphiphilic peptide reagent to mediate intracellular delivery of the same pre-formed CRISPR ribonucleoprotein enzymes that are broadly used in research and therapeutics, resulting in high-efficiency editing of stimulated immune cells and cultured hematopoietic stem and progenitor cells (HSPCs). PERC facilitates nuclease-mediated gene knockout, precise transgene knock-in, and base editing. PERC involves mixing the CRISPR ribonucleoprotein enzyme with peptide and then incubating the formulation with cultured cells. For efficient transgene knock-in, adeno-associated virus (AAV) bearing homology-directed repair template DNA may be included. In contrast to electroporation, PERC is appealing as it requires no dedicated hardware and has less impact on cell phenotype and viability. Due to the gentle nature of PERC, delivery can be performed multiple times without substantial impact to cell health or phenotype. Here we report methods for improved PERC-mediated editing of T cells as well as novel methods for PERC-mediated editing of HSPCs, including knockout and precise knock-in. Editing efficiencies can surpass 90% using either Cas9 or Cas12a in primary T cells or HSPCs. Because PERC calls for only three readily available reagents – protein, RNA, and peptide – and does not require dedicated hardware for any step, PERC demands no special expertise and is exceptionally straightforward to adopt. The inherent compatibility of PERC with established cell engineering pipelines makes this approach appealing for rapid deployment in research and clinical settings.

## INTRODUCTION

CRISPR-mediated genome editing has rapidly revolutionized biological research and enabled several clinical trials evaluating the technology’s therapeutic potential. There are many cell, tissue, and organ contexts in which *in vivo* genome editing could address substantial unmet medical need, however limitations associated with delivery technology have thus far prevented safe and effective use of candidate CRISPR therapies outside of the liver^1^. Fortuitously, transplantation of certain cell types allows CRISPR-mediated cell therapies to be engineered in the laboratory setting and administered to patients via *ex vivo* therapy. Established transplantation regimens have allowed for incorporation of CRISPR into trusted cell therapy pipelines, largely sidestepping delivery challenges and facilitating clinical assessment of potent genome editing technologies. For example, autologous hematopoietic stem and progenitor cell (HSPC) transplantation, which is widespread in the treatment of blood cancers, was adapted to incorporate CRISPR, enabling several trials to address hemoglobinopathies and resulting in the first approved CRISPR therapy^2^. Widespread viral engineering of T cells *ex vivo* to generate anti-cancer cell therapies hastened the adoption of CRISPR-enhanced versions, leading to academic clinical trials^3,4^ and several companies developing immuno-oncology approaches including autologous (“off-the-shelf”) T cell therapies^5–7^. *Ex vivo* genome editing of immune cells and HSPCs also offers hope for the treatment of primary immunodeficiencies, which are often severe conditions with immense unmet need. Beyond these encouraging clinical applications, CRISPR-mediated editing of T cells and HSPCs is an area of fundamental research. CRISPR screening in T cells has revealed a wealth of functional information^8,9^, and screening in HSPCs has been used to model clonal hematopoiesis^10^ or to identify strategies for addressing hemoglobinopathies^11,12^.

Even with the ease of CRISPR delivery in the laboratory context, T cells and HSPCs are relatively fragile and generally resistant to transfection, leading researchers and clinical development teams alike to rely on electroporation of CRISPR enzymes, typically in ribonucleoprotein (RNP) format, as the predominant delivery technology for precise engineering of these cells. Electroporation generally promotes high editing efficiencies but is associated with substantial drawbacks: electric current can dramatically perturb or kill cells, and it necessitates both dedicated hardware and cumbersome transport of cells to and from the reaction device. To avoid these limitations while retaining high editing efficiency in T cells and HSPCs, we describe the development and recommended methods for use of peptide-enabled RNP delivery for CRISPR engineering (PERC).

### Development of PERC

Delivery of CRISPR enzymes into immune cells or HSPCs using the RNP format has many advantages: both protein and RNA can be readily manufactured, the assembly step is simple and effective, the resulting enzyme is immediately active in the cell, and its short half-life minimizes off-target editing^13^. Intracellular delivery of CRISPR RNP enzyme is usually carried out by electroporation, which can result in efficient editing but comes with significant complications. The field has lacked a gentle, convenient, potent, reagent-based delivery technology for clinically important cells in blood and immune system disorders. Recently, the use of amphiphilic peptides as a delivery method for CRISPR RNPs has been reported by several groups^14–19^. PERC is a straightforward, relatively inexpensive, and efficient method for delivery of CRISPR RNPs in immune cells, does not require specialized hardware, and is gentler than electroporation^15,16^ (**Fig. 1a**).

**Figure 1.**
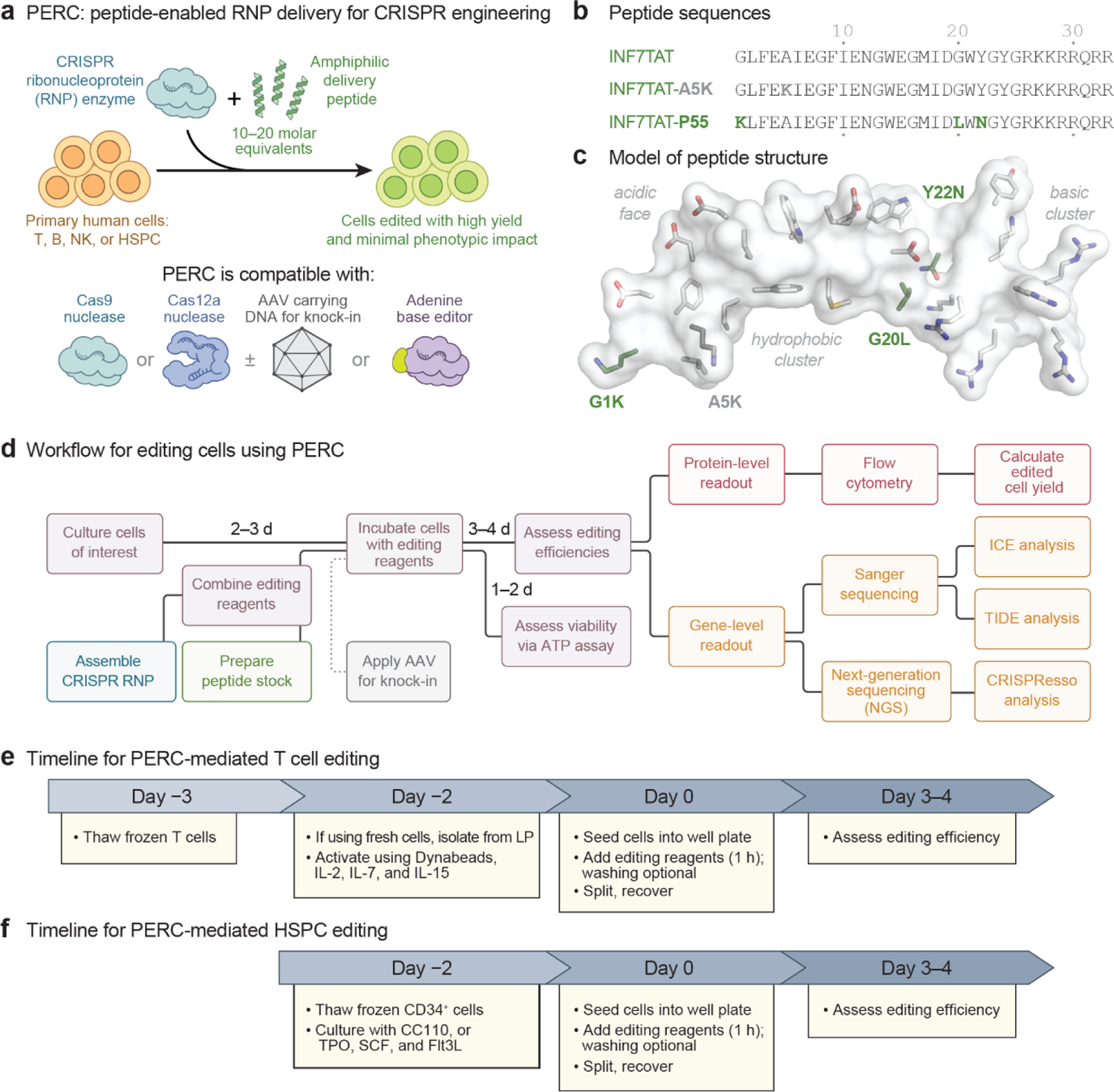
PERC for editing T cells and HSPCs. **a.** Overview of PERC, depicting the reagent-based approach for CRISPR enzyme delivery to primary human cells. CRISPR RNP is combined with PERC peptide and co-incubated with cultured immune cells to achieve genome editing. PERC has been used with base editors as well as Cas9 or Cas12a nuclease; the nucleases can be paired with AAV HDRT to facilitate transgene knock-in. **b.** The sequence of INF7TAT and the derived peptides A5K and P55, either of which can enable PERC. Amino acid substitutions that confer improved activity are in bold. **c.** A space-filling structural model of the INF7TAT peptide scaffold. The INF7 portion is based on the solution structure of HA2 (PDB ID: 1IBN) and the TAT portion is modeled as an idealized alpha helix. Oxygen and nitrogen are represented as red and blue, respectively, and the carbon color varies. Key residues are represented as sticks (for side chain atoms beyond the α-carbon), and light gray represents the invariant hydrophobic, acidic, and basic residues. The A5K substitution is in dark gray, and the trio of P55 substitutions are in green. Note that the A5K substitution is not present in P55. Model created using PyMOL (v1.1r1). **d.** Workflow of the steps and timing for PERC. The optional use of AAV to support efficient HDR-mediated knock-in is represented by a dotted line. **e.** Timeline of primary human T cell culture with PERC. LP refers to leukopak. **f.** Timeline of primary human HSPC culture with PERC.

We previously used PERC to perform efficient genome editing in primary human T cells while preserving a naïve cell phenotype, minimally perturbing the transcriptome, minimizing chromosomal translocations, generating a higher yield of precisely integrated chimeric antigen receptor (CAR)-T cells than with electroporation, and establishing compatibility with multiple rounds of delivery. PERC was compatible with serial delivery for multiplex editing, resulting in knockout at three loci and knock-in of a CAR gene using an AAV6 homology-directed repair (HDR) template (HDRT). Sequential multiplex editing also supported double knock-ins, and the production of CAR-T cells that displayed potency in a mouse model of B-cell leukemia^15^.

We also previously demonstrated efficient PERC-mediated genome editing of T cells using Cas9, Cas12a, and an adenine base editor, as well as Cas9-mediated editing of B cells and NK cells. These experiments used the A5K (alanine to lysine) substitution variant of the INF7TAT peptide^15^. INF7TAT peptide had served as the basis for a prior screen for lytic activity in red blood cells as a proxy for endosomal escape^20^. Our screen of INF7TAT variants in T cells^15^ identified A5K as well as three additional activating INF7TAT substitutions (G1K, G20L and Y22N) that we have now incorporated in a single peptide: INF7TAT-P55 (henceforth P55) (**Fig. 1b,c**). Like the A5K variant, P55 supports efficient genome editing in T cells with excellent yields, and it is well-suited for CD34^+^ HSPCs. Both A5K and P55 support PERC in HSPCs, and the latter is used throughout this work because it promotes slightly improved editing rates in these cells.

In this manuscript, we elaborate a detailed, reproducible method to carry out PERC in primary human T cells and HSPCs. We have optimized PERC doses for both cell types to achieve higher editing efficiency and – in alignment with the recently reported “PAGE” method for peptide-mediated RNP delivery^16^ – incorporated an abbreviated incubation time followed by washing to remove PERC reagents. This modification can maintain editing efficiency while improving cell viability. We also describe steps for PERC-mediated delivery of CRISPR RNP paired with AAV HDRT to facilitate transgene knock-in for T cells and HSPCs. We previously demonstrated efficient knock-in for T cells^15^, and now report an approach that works for HSPCs as well. PERC-mediated knock-in for HSPCs produces editing efficiencies comparable to those attained with electroporation or lipid nanoparticles^21–24^, with the added advantages of PERC.

### Applications of PERC

Using PERC, we have shown efficient delivery of Cas9, Cas12a, and an adenine base editor in immune cells^15^. We anticipate that PERC would also support delivery of other enzymes in the genome editing toolkit such as other Cas12 enzymes, Cas13 enzymes, epigenome editors, prime editors, cytosine base editors, and non-CRISPR enzymes such as zinc finger nucleases, meganucleases, or recombinases. While it remains to be explored, PERC may prove useful in genome editing of non-immune or hematopoietic cells, especially those that are recalcitrant to transfection or too delicate to endure electroporation.

PERC is gentle on cells, permitting sequential editing of multiple loci. As previously reported, this is one way to minimize chromosomal translocations^15,25^. This approach has resulted in near-complete elimination of translocations with spacing of 2 d^15^ or 3 d^25^ between instances of nuclease delivery. Spacing PERC delivery steps by ≥ 2 d allows each RNP to be metabolized by the cell^26^, in turn ensuring that the double-strand break (DSB) at each locus is resolved, which limits the opportunity for translocation between DSBs at multiple loci^15^. This benefit might contribute to precise multiplexing in cell therapy products without incurring the risk of translocations, which are a cause for clinical concern^27^. Because PERC is minimally toxic and calls for no dedicated hardware, we anticipate that PERC could conveniently be paired with other modes of delivery (e.g. electroporation, AAV, lentivirus, or lipid nanoparticles) to support engineering of sophisticated cell therapy candidates that have previously been impossible or impractical due to delivery constraints. Furthermore, a similar multi-step editing strategy could facilitate hypothesis-driven research investigating complex genetic interactions: a gene of interest could be knocked out using PERC in primary cells, and a second editing step could use a traditional pooled screen to examine the role of other genes in the knockout context.

### Comparison with other delivery methods

T cell engineering has predominantly relied on gammaretroviral or lentiviral vectors for transgene delivery. Despite high transduction efficiency in both dividing and non-dividing cells, viral vectors produce heterogeneous cell populations with lower transgene activity than that of cells with precise engineering for homogeneous transgene expression^30,31^. Viral transduction is flexible in terms of hardware requirements, and there is established global infrastructure that supports viral transduction of T cells for therapeutic use. Despite the technical advantages offered by precise CRISPR engineering of cell therapies, widespread adoption has been slow, in part due to the practical hurdles associated with incorporating electroporation into manufacturing pipelines. We imagine that PERC could help bridge this gap, facilitating precise CRISPR engineering in the same manufacturing context that has been established for viral transduction.

Among non-viral methods, electroporation and mRNA/LNP-based methods have been most successful for CRISPR enzyme delivery into immune cells and HSPCs. Physical methods such as electroporation^32^ and microfluidic delivery^33^ can produce high editing efficiencies. Electroporation has a track record as a clinically effective means of delivery, enabling multiple clinical trials (NCT03745287, NCT05942599, NCT04637763, NCT05757700) and the world’s first CRISPR-edited cell therapy, Casgevy^34^. While electroporation promotes efficient delivery and editing, it comes at the cost of cell health and viability^24,33,35^. Edited cell yield and condition is of immense importance in allogeneic T cell therapies, which ideally produce as many effective doses as possible per manufacturing run^36^, and when editing the fragile and scarce HSPCs underlying autologous cell therapy for sickle cell disease^37^. Peptide-mediated RNP delivery^15,16^ could be an effective alternative delivery method, streamline manufacturing pipelines, and be compatible with “closed loop” manufacturing as well as straightforward incorporation into established hardware such as the CliniMACS Prodigy or G-Rex devices^38,39^.

LNPs can co-deliver CRISPR guide RNA (gRNA) and protein-encoding mRNA in a hardware-independent fashion, and, like PERC, can be gentler than electroporation^24^. LNPs have only recently emerged as an option for *ex vivo* manufacturing of T cell or HSPC therapies and have not yet reached the clinic for these cell types. LNPs hold substantial promise for *in vivo* use in T cells and HSPCs, having demonstrated clinical success in CRISPR delivery to the liver^40^ and recently showing promise for *in vivo* T cell delivery in preclinical models^41,42^. Furthermore, the feasibility of LNP manufacturing at massive scale was established by the COVID vaccines^43,44^. However, LNPs require reagents that call for complicated synthesis and exacting storage conditions. Alternatively, they can be purchased from vendors in kit format, which is substantially more expensive than a synthetic delivery peptide. In comparison to making (or purchasing) the protein and gRNA components of RNP, synthesis of high-quality mRNA can be expensive and cumbersome^45^. These practical considerations may underlie the slow adoption of LNPs for *ex vivo* CRISPR delivery. Optimal preparation of LNPs require a microfluidic instrument, which involves substantial dead volumes and associated material loss, and is ill-suited to generating numerous samples in a medium- or high-throughput scenario^46^. In contrast, PERC can be readily performed at small scale, enabling rapid screening of various editing reagents via parallel or serial delivery. PERC is also extremely convenient, calling for simple combination of an inexpensive peptide and the CRISPR RNP, a cargo format already in widespread use for electroporation. Finally, PERC reagents are generally stable, with minimally demanding storage conditions and compatibility with freeze-thaws.

### Expertise needed to use PERC

CRISPR delivery into T cells or HSPCs via PERC requires no special expertise. As described below, procuring the reagents for peptide-mediated RNP delivery is straightforward if one relies upon commercial vendors. Alternatively, labs with appropriate capacity and experience may opt to generate any of the three core components: recombinant CRISPR protein can be expressed in *E. coli* and purified using standard chromatography methods^47^; high-purity gRNA can be produced via *in vitro* transcription followed by PAGE purification^47^; and amphiphilic peptide can be obtained via solid-state synthesis followed by HPLC purification^48^. For AAV HDRT knock-in, in-house virus production may be preferable as commercial production can be expensive.

Post-PERC editing efficiency can be evaluated in several ways: flow cytometry offers a rapid protein-level assessment of gene expression; Sanger sequencing provides convenient low-resolution quantification of genomic changes; and amplicon-based next-generation sequencing (NGS) offers the most accurate readout of editing at a locus. For NGS analysis of editing outcomes, some coding experience can be beneficial but is not necessary. Both web-based and more complex command line toolsets are freely available.^49^

### Selecting and procuring guide RNA (gRNA or crRNA)

Guide RNA can be designed by following guidelines described previously^50^ for using tools such as CHOPCHOP, CRISPOR, etc. or by selecting a guide that has been used in previous studies. We recommend testing several guides to identify one that works best. In general, guides that are extremely efficient via electroporation of RNP also tend to excel via PERC. We purchase guides from Integrated DNA Technologies (IDT), Synthego, or Horizon Discovery with the manufacturer-recommended standard chemical modifications. Introducing chemical modifications to the gRNA has been shown to markedly improve the potency of resulting RNPs in at least one use case^51^, but this is not universally true^52,53^. We note the potential for comparable activity because one can affordably produce a guide lacking chemical modifications using *in vitro* transcription, as described previously^47^, ideally with removal of the triphosphate cap.^54^ While improving gRNA purity has not been definitively demonstrated to improve editing efficiency, gRNA heterogeneity can be largely eliminated by flanking the transcript with ribozymes that leave precise 5′ and 3′ gRNA termini^17,55^, which also removes the immunogenic triphosphate.

Single guide RNA (sgRNA) and dual guide RNA (dgRNA; comprising a separate tracrRNA and crRNA) tend to work interchangeably, and both are compatible with PERC. For optimal activity, the use of dgRNA may call for an increased RNA:protein ratio as compared to sgRNA^23^. The appeal of dgRNA is that it allows for purchase of a large quantity of tracrRNA, which is not target specific, and smaller amounts of target-specific crRNA, which are shorter in length and hence easier, faster, and cheaper to synthesize. This strategy may be efficient when screening many targets.

### Selecting and procuring Cas protein

It is essential to select the right type of CRISPR effector for a specific need. This choice depends on the genomic locus that one wishes to edit, since CRISPR nucleases require a protospacer-adjacent motif (PAM) adjacent to the intended cut site, e.g. NGG for SpCas9; NNGRRT for SaCas9; TTTV for many Cas12a enzymes. Despite the widespread use of SpCas9, we recommend comparing it to Cas12a at the outset of a project, as the latter enzyme often excels via PERC delivery.

For using SpCas9, several different constructs are available, and some work especially well with PERC. We recommend the Cas9-triNLS construct (referred to as “3xNLS” when initially published), which has three different nuclear localization signals: SV40, c-Myc, and nucleoplasmin^56,57^. This protein tends to purify with high yield and works well with PERC, making it our top pick for non-industrial use^15^. To our knowledge Cas9-triNLS is not commercially available but can be requested for purchase from MacroLab (a core at University of California, Berkeley). It is also available as expression plasmids for purification using previously reported methods (Addgene #196244 and #196245)^47^ or can be synthesized by a contract research organization such as Aldevron^58^. Another potent option is ‘Cas9-6xNLS’ (initially referred to as “4xNLS”), which contains six SV40 NLS sequences (4 N-terminal & 2 C-terminal) that promote moderate levels of RNP self-delivery into neurons (Addgene #88917 and #196246)^59^. Although we primarily used this protein in our previous work^15^, it can be challenging to express and purify in high yield, so we do not recommend it. Of the commercially available SpCas9 proteins that we evaluated, TrueCut V2 protein performed the best with PERC^15^. The recently reported Cas9-T6N construct contains an N-terminal TAT CPP fusion with four C-Myc NLS as well as two C-terminal SV40 NLS (Addgene #199604); this protein worked well via peptide-mediated RNP delivery^16^ and we anticipate it would work well with the PERC as described herein.

For Cas12a, our preliminary evaluation for PERC suggested that AsCas12a outperformed LbCas12a (unpublished data). We proceeded with the Alt-R AsCas12a Ultra enzyme (IDT), which contains point mutations that enhance enzyme activity without sacrificing specificity and works well with PERC^15,60^. This protein contains a single SV40 NLS, and previous work with Cas9^15,56,57^ and Cas12a^16,61^ suggested that increasing NLS density can enhance activity. We evaluated two AsCas12a constructs: Cas12a-Ultra-2xNLS with two SV40 NLS sequences fused to the C-terminus, and a Cas12a-Ultra-5xNLS with an SV40 sequence fused to each terminus plus C-terminal fusion of three more NLS (nucleoplasmin and a pair of C-Myc). The two NLS-enriched constructs performed similarly in T cells via PERC, while Cas12a-Ultra-5xNLS supported markedly improved PERC-mediated edited cell yield in HSPCs. Here we used Cas12a-Ultra-2xNLS for T cells experiments and Cas12a-Ultra-5xNLS for HSPC experiments. Cas12a-Ultra-5xNLS is available as a plasmid for expression in *E. coli* (Addgene #218775), and both constructs are available as ready-to-ship protein to purchase from MacroLab by request. Another recently reported construct, opCas12a-T8N (Addgene #199605), which is based on enAsCas12a^62^, has six N-terminal c-Myc NLS sequences plus two C-terminal SV40s sequences and supports efficient editing via peptide-mediated delivery^16^.

### Procuring peptide for PERC

Two PERC peptides are commercially available: INF7TAT-A5K (A5K)^15^ and INF7TAT-P55 (P55). Both peptides are compatible with PERC, but P55 tends to have slightly higher potency in T cells and HSPCs. Peptide (sequences in **Fig. 1b**) can be synthesized either by a vendor or in the laboratory using established protocols^48,63^. For synthesis, we recommend ≥95% purity as assessed by HPLC, which is also used for purification. It has been suggested that an acid exchange step (using HCl or acetate to displace trifluoroacetic acid) can support cell health^16^, but in our experience this step may not be a critical consideration.

### AAV6 HDRT design (optional)

For gene knock-in, we designed an AAV-delivered homology directed repair template (HDRT), which is the exogenous DNA sequence to be integrated at the site of nuclease-induced DSB. This reagent is designed to enforce knockout of *B2M* and introduce a transgene. The transfer plasmid, which incorporates the HDRT sequence into the AAV6 vector, contains the following sequence elements: a first inverted terminal repeat; left homology arm corresponding to upstream of the gRNA cut site (in *B2M* exon 2); P2A ribosome skipping sequence; truncated nerve growth factor receptor gene containing a signal peptide, ectodomain, and transmembrane domain; stop codon and bGH polyA tail; right homology arm corresponding to downstream of the gRNA cut site; and a second inverted terminal repeat. Design principles and best practices for AAV HDRT delivery have been detailed elsewhere.^21,30^

### Procuring AAV6 (optional)

We use recombinant AAV6 to deliver the HDRT. Commercial vendors offer AAV synthesis for research or clinical use. Alternatively, AAV can be produced using established protocols^64–66^. Here, we made AAV by transfecting HEK293T cells with cargo plasmid, rep-cap plasmid, and helper plasmid using PEI, and then purified AAV using density gradient ultracentrifugation. The titer was estimated using qPCR as the concentration of viral genomes (vg/µl), and virus was aliquoted and stored at −80°C.

### Procuring ssODN to facilitate HDR (optional)

T cell editing by PERC to knock in short sequences can be achieved using a single-stranded oligodeoxynucleotide (ssODN) as the HDRT, which can be commercially synthesized by vendors such as IDT. We found that the ssODN HDRT design crucially required incorporation of a truncated Cas9 target site on the HDRT to enhance RNP interactions^23^. Additional design parameters have been described previously^67,68^.

### Assessing editing efficiency

Genome editing outcomes can be quantified via several assays, some of which depend on the target locus (**Fig. 1d**). Changes in protein expression to evaluate knockout or knock-in can be assessed by flow cytometry or western blotting. Flow cytometry is particularly useful in the evaluation of engineered T cells and HSPCs because their surface properties are often of great importance, and cell phenotype can be simultaneously evaluated. For some proteins, knockout is apparent only after biallelic gene disruption. Because monoallelic edits often go undetected by flow cytometry (or blotting), apparent editing rates may seem to differ when comparing protein-level assays to DNA-level assays that assess individual alleles. In addition to directly assessing editing efficiency at the DNA level, sequencing can reveal the precise details of the indel profile (the sequence changes at the targeted locus). Amplicon-based NGS is a powerful and sensitive method that provides accurate data on editing at the DNA level but may involve partnering with a core service that is expensive. NGS analysis can be carried out using tools such as CRISPResso^49^ (http://crispresso2.pinellolab.org/submission), which can be used manually for a small number of samples. For larger datasets, however, it may be useful to have some scripting background. We use Cortado (https://github.com/staciawyman/cortado), which is a reimplementation of CRISPResso 1.0 that offers users the ability to analyze hundreds of samples with ease. An alternative option for DNA editing analysis is Sanger sequencing, which is typically much faster and less costly but does not provide sequence information for individual edited alleles, can under-report editing rates, and struggles to detect editing when rates are ≤ 10%. Sanger sequencing data can also be analyzed using free online tools such as ICE (Inference of CRISPR edits)^69^ or TIDE/TIDER (Tracking of Indels by Decomposition)^70,71^. For analyzing base editing outcomes, flow cytometry can be used for edits that alter protein expression, while sequence-level readout can be provided by Sanger-based analysis^72^ and/or amplicon-based NGS. Here we provide DNA-level data on indel-mediated knockout via both amplicon-based NGS and Sanger sequencing.

### Limitations

Any transient delivery method, be it PERC, mRNA/gRNA-bearing LNP, or electroporation of RNP or mRNA/gRNA is unlikely to attain the same editing efficiencies resulting from long-term expression of editing machinery. Although electroporation of RNP typically surpasses PERC in terms of absolute editing efficiencies, improvements in viability often cause the yield of edited cells to be higher following PERC^15^. PERC is optimized for actively dividing T cells and HSPCs. This aspect may be a limitation, as recent studies have shown that editing resting cells can reduce chromosome fragment loss in both T cells^73^ and HSPCs^74^. That said, it is possible that the genotoxicity observed in those studies is rooted in electroporation itself and is exacerbated by the process of cell division. Further study will be needed to determine whether genotoxicity is attenuated with PERC in cycling cells, and whether efficient PERC is possible in resting cells.

For knock-ins, single-digit percentage efficiency HDR via PERC is possible with peptide-mediated delivery of short ssODN HDRTs, which are readily commercially available. While this has not been tested in HSPCs, we previously reported these results in T cells^15^. High-efficiency HDR via PERC can be achieved by using AAV6 for HDRT delivery. The use of AAV in conjunction with electroporation-based delivery of CRISPR RNP has been shown to impair the capacity for HSPCs to engraft^22^ and might have contributed to complications in a recent clinical trial^75^. However, it remains to be seen whether these consequences can be alleviated in the absence of electroporation, a harsh process that could potentially synergize with AAV to impair HSPC engraftment potential. In light of this unknown, the use of PERC for CRISPR RNP delivery may be an appealing alternative for further investigation.

### Experimental design

#### Cell culture

It is important to plan for and set aside a reasonable amount of time for culturing cells several days in advance of the day when cells will be edited (**Fig. 1e,f**). Primary cells may support higher editing efficiencies when they are actively dividing, so it can be beneficial to introduce the editing reagents at the appropriate time. For primary human T cells, we recommend activating cells two days before editing. To activate cells on Day −2 (relative to the day of editing), we recommend thawing frozen T cells three days prior to editing, and letting the cells rest and recover overnight from the thaw. If T cells will be isolated from a leukopak and used immediately for an experiment, this can be done two days prior to editing (**Fig. 1e**). We have compared freshly isolated T cells and previously frozen T cells and observed no substantial difference in editing efficacy or viability. For HSPCs, we recommend thawing frozen cells two prior to editing (**Fig. 1f**).

#### Design of an initial PERC experiment and controls

We suggest initially testing a range of doses to identify an optimal dose. We recommend including one condition that contains RNP without peptide and another condition that contains peptide without RNP. These conditions should result in low or no editing and serve as negative controls. The peptide-only condition can be used to test a given cell type’s sensitivity to the peptide, although we note that peptide-mediated toxicity can be exacerbated by the absence of RNP cargo. Non-treated negative controls should be included to establish baseline properties (e.g. target protein expression, cell health). For a positive control, we suggest using a method such as electroporation if the necessary hardware is available. Dedicated protocols for electroporation of T cells^32,76^ and HSPCs^21^ are available, and peptide-mediated RNP delivery has been extensively compared with electroporation in previous studies^15,16^.

Below is an example 96 well plate experiment with a treatment volume of 100 µL, for an initial dose optimization, including controls:

- 100 pmol RNP + 1 nmol peptide (10 μM peptide in 100 μL)
- 200 pmol RNP + 2 nmol peptide (20 μM peptide in 100 μL)
- 400 pmol RNP + 4 nmol peptide (40 μM peptide in 100 μL)
- 0 pmol RNP + 4 nmol peptide (40 µM peptide in 100 μL) (replace RNP volume with equal volume of RNP buffer)
- 400 pmol RNP + 0 nmol peptide (replace peptide stock with equal volume of DMSO)
- NT (Non-treated) 0 pmol RNP + 0 nmol peptide (replace RNP volume with equal volume of RNP buffer and replace peptide stock with equal volume of DMSO)

Calculate the volume of gRNA and Cas protein needed based on the total amount of RNP required for the experiment. For the above example doses, 1.1 nanomoles of RNP are needed. We suggest making 20% extra to account for pipetting loss, and enough for two replicates, which would be 2.64 nanomoles of RNP. This example maintains a molar ratio of 10:1 for peptide:RNP, though testing other ratios such as 5:1 and 20:1 may be informative. Various peptide:RNP ratios can be evaluated, keeping fixed either the amount of RNP or the amount of peptide. Testing different concentrations of both reagents in a 2×2 matrix can help identify an optimal dose and ratio of the key reagents.

It may also be useful to determine the optimal ratio of gRNA:protein for the RNP assembly step. While the field has reached a general consensus that a molar excess of gRNA is beneficial for efficient RNP assembly and editing activity^23^, we note that a given enzyme, gRNA format (sgRNA, dgRNA, or crRNA), and/or target-specific spacer sequence might perform best at different gRNA:protein ratios. In our experience, the optimal gRNA:protein ratio for a given target-specific nuclease is independent of the method for RNP delivery. For this reason, a ratio that has been determined via electroporation need not be reoptimized for PERC. If empirically determining this ratio for the first time, we recommend comparing gRNA:protein molar ratios of 1.5, 2, 3, and 4. A gRNA:protein ratio of 1.5 is optimal for many target-specific RNPs, so we recommend this as a starting point.

For knock-in experiments, control conditions can be included for knockout editing (RNP and peptide, without AAV) and for AAV alone (without RNP or peptide). Optimization may be needed to identify an appropriate multiplicity of infection (MOI) for AAV transduction. For T cells, we find that 5×10^4^ vg/cell typically appropriately balances the objectives of obtaining high knock-in efficiency and not using up excess AAV reagent; higher MOIs provide diminishing returns on knock-in efficiency. For HSPCs, an additional objective is to limit the MOI to preserve cell health, such that a lower amount (e.g. 2.5×10^4^ vg/cell) may be more appropriate.

#### Assessing editing efficiency

To assess editing efficiency, it is important to consider the time needed for the target protein’s turnover. This is relevant when selecting a timepoint for assessing protein expression and may require optimization. In the present work, we have targeted beta-2-microglobulin (*B2M*), which is expressed on almost all human cell types and is pertinent to allogeneic immune cell therapies^77^. B2M knockout can be assessed by flow cytometry three to four days after delivery of editing reagents. This time is sufficient to distinguish expressing and knockout cells, with four days improving the distinction between positive and negative populations by flow cytometry.

#### Designing a panel for flow cytometry

We recommend designing a flow cytometry antibody panel for the target of interest plus cell-identifying markers using tools such as the Thermo Fisher flow panel builder (www.thermofisher.com/order/panel-builder/). For T cells, we recommend staining for CD3, CD4, and CD8. A viability dye such as Ghost Dye Red 780 or 7-AAD distinguishes live and dead cells. For HSPCs, we recommend staining for CD34 as a classic stem and progenitor cell marker.

#### Designing primers for sequencing

To assess editing efficiency by sequencing, sequencer-specific requirements should be followed. Designing primers that sufficiently flank the intended cut site is critical to observe the spectrum of possible insertions or deletions (indels). For example, to perform 300 bp paired-end Illumina sequencing, we made an amplicon around 300–400 bp and used the MiSeqx300 kit. For Sanger sequencing, we recommend an amplicon of 600–1,000 bp. Primers can be designed using tools such as Primer3^78–80^, specificity can be assessed by Prime-rBLAST^81^. Table 3 provides a list of primers designed for the *B2M* gRNAs in Table 2.

**Table 1:**
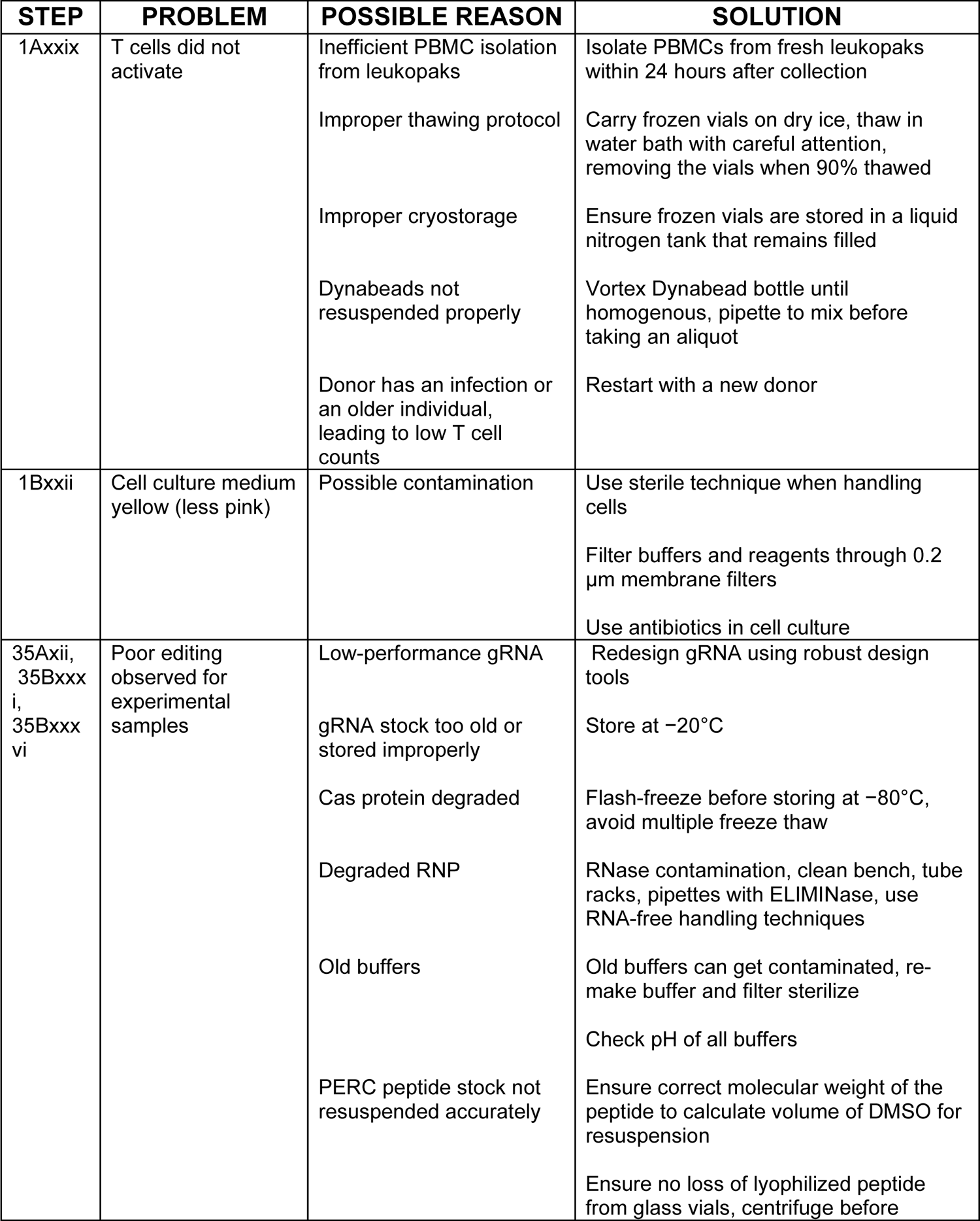
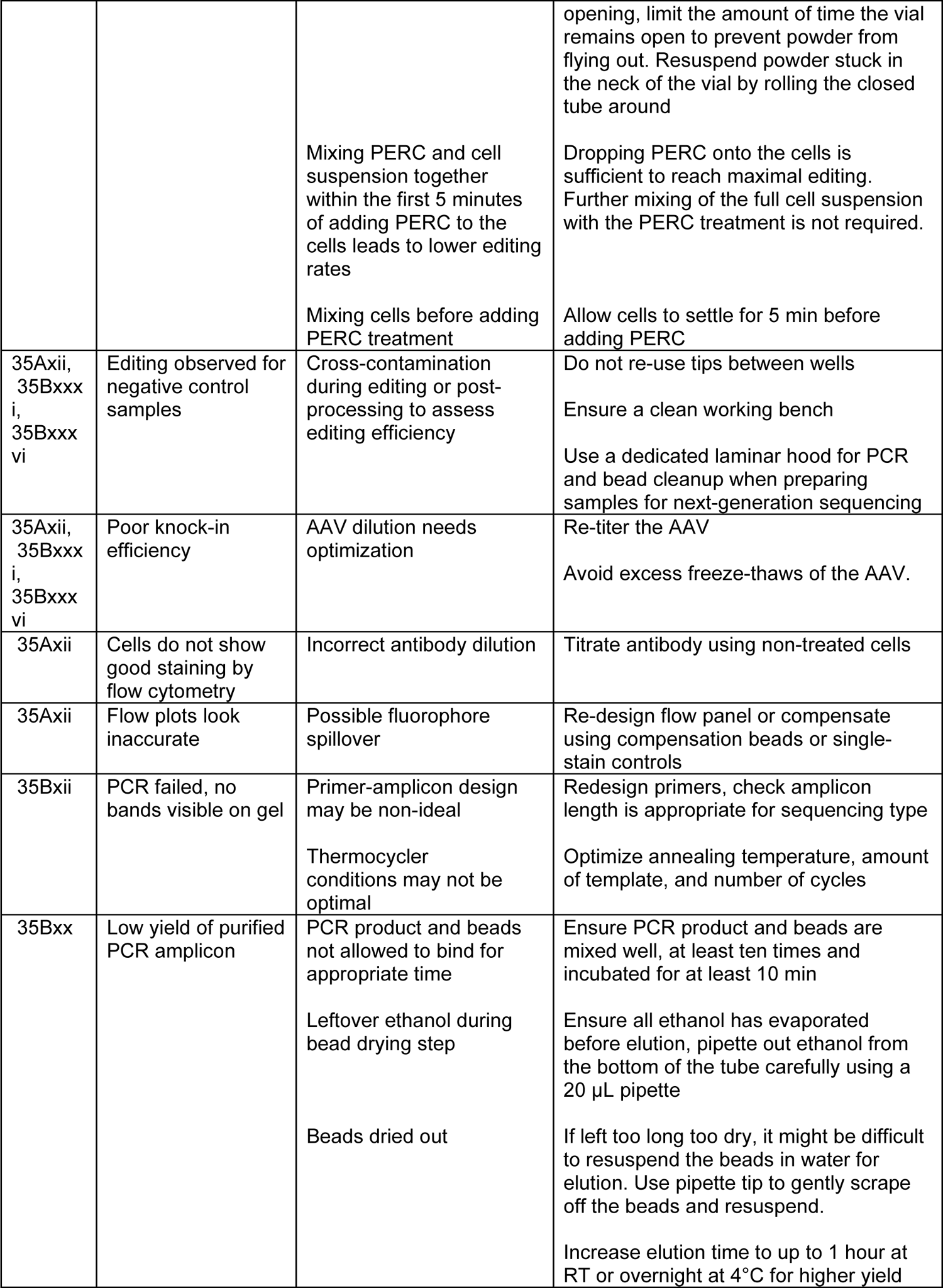
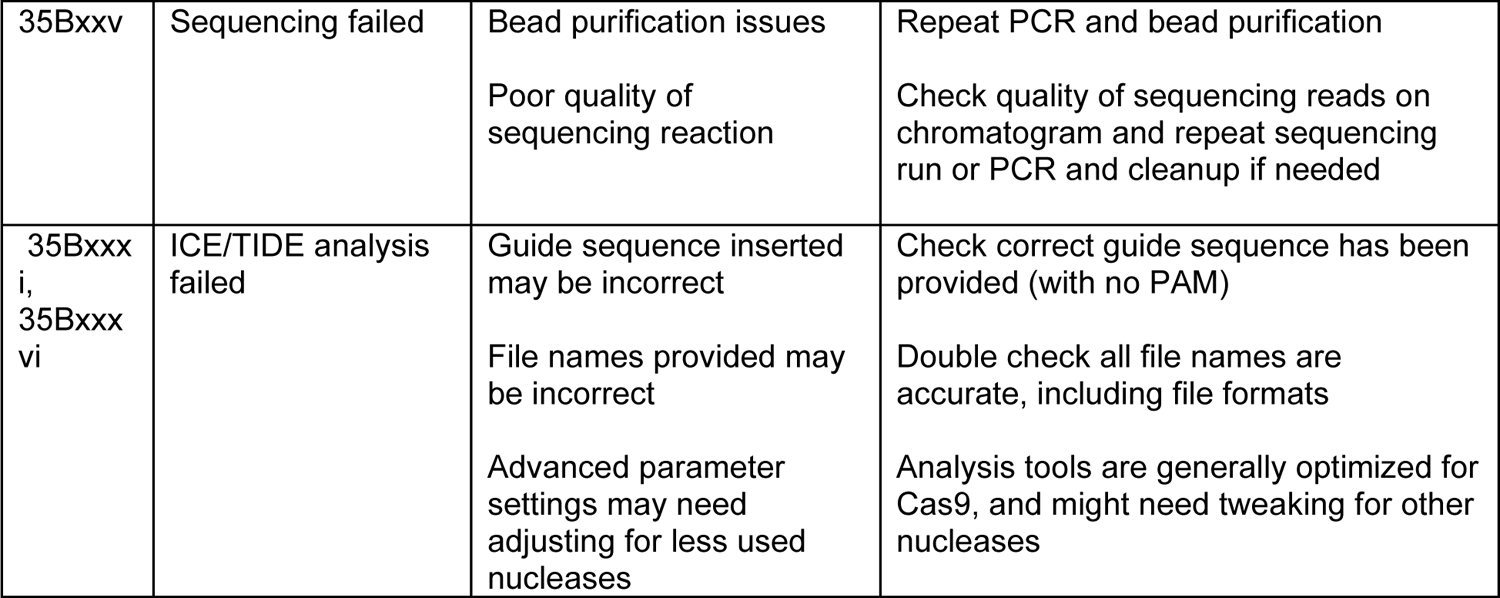
Troubleshooting table. Table listing potential problems that may arise when using PERC, with recommended troubleshooting steps.

**Table 2:**
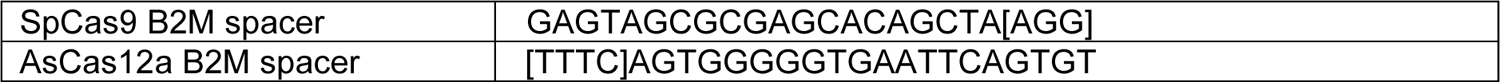
gRNA sequences. Table listing all guide RNA sequences used. Protospacer Adjacent Motif (PAM) sequences are shown within brackets.

**Table 3:**
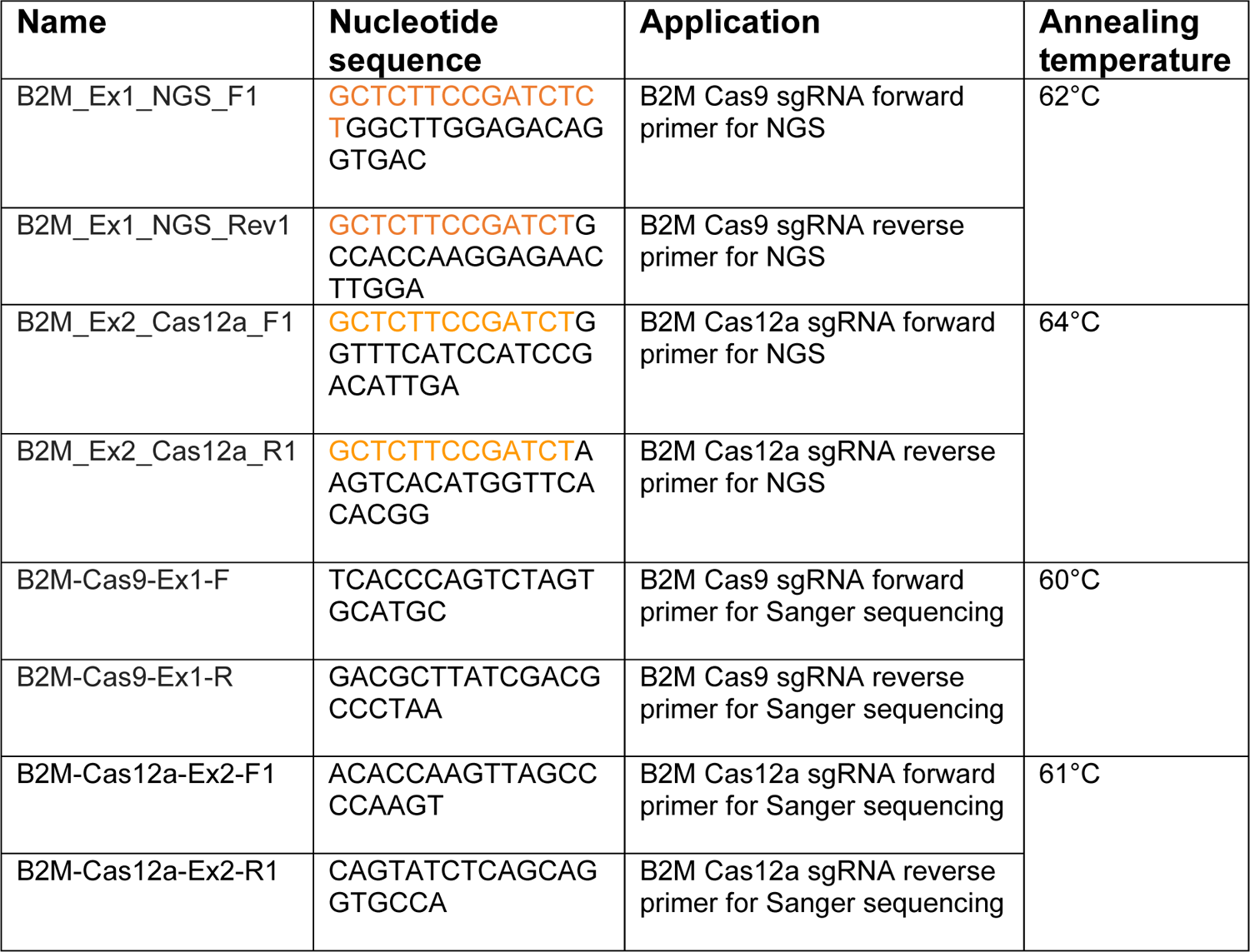
Oligo sequences. List of all oligonucleotides used throughout. Nucleotides shown in color are stubs used for the second PCR amplification for NGS.

## RESULTS

PERC is an efficient and convenient method for delivery of CRISPR in the form of RNPs into T cells and HSPCs. Using PERC, we routinely achieve 50–80% protein-level knockout with SpCas9 and 80–95% with AsCas12a across different loci. When paired with an AAV6 HDRT, we routinely achieve 45–55% knock-in for T cells and 15–40% knock-in for HSPCs.

**Fig. 2** and **Fig. 3** show results from the PERC-mediated delivery of CRISPR enzymes targeting the *B2M* locus in primary human T cells using SpCas9 RNP or AsCas12a RNP, respectively. Rates of cell-surface protein knockout as assessed by flow cytometry should be in the range of 50–80% when using Cas9, and 75–95% when using Cas12a. Editing efficiencies assessed using NGS provide slightly different values because sequencing detects alleles independently, while flow cytometry assesses protein knockout resulting from biallelic editing in a given cell. Although Sanger sequencing results tend to agree with NGS when editing rates are close to 100%, Sanger sequencing tends to under-report editing as efficiency decreases. **Fig. 4** and **Fig. 5** show results for analogous experiments in HSPCs, where flow cytometry detects 60–75% knockout via Cas9 and 85–95% knockout via Cas12a. Generally, increasing the dose of RNP and peptide leads to higher editing efficiencies, although cell viability may decrease at higher doses. Washing off the treatment after 1 hour of incubation may help improve cell health, and hence survival and edited cell yield. **Fig. 6** shows representative results when PERC is paired with an AAV HDRT for gene knock-in in HSPCs: ∼30% precise knock-in, with ∼80% of cells being edited (including knock-out or knock-in). PERC does not require specialized hardware and can be readily applied to make multiplex-edited CAR-T cells, to correct pathogenic alleles in HSPCs, or to introduce myriad other edits in these primary cells.

**Figure 2.**
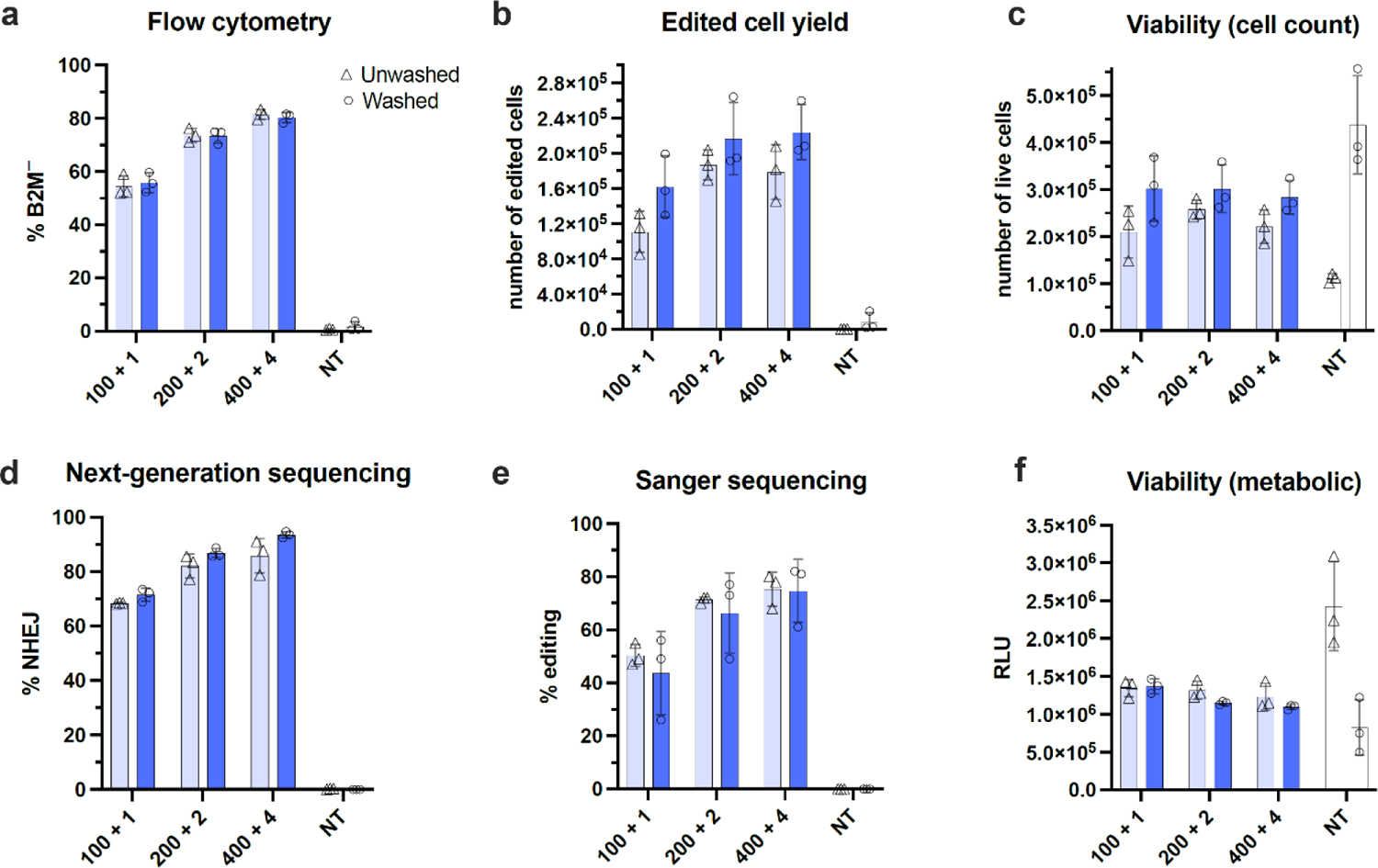
Cas9-mediated editing of primary human T cells using PERC. Primary human T cells were incubated with editing reagents for 1 hour in Opti-MEM followed by either a 4× dilution (unwashed treatment) or washing off editing reagents (washed treatment) and recovering cells in growth medium supplemented with 5% FBS and 300 U/mL IL-2. **a.** Protein-level KO of B2M assayed using flow cytometry. **b.** Number of cells with knockout of B2M surface expression, as assayed in (a). **c.** Cell viability depicted as number of live cells determined by Ghost Dye Red 780 staining four days after editing. **d.** DNA-level editing assessed by Illumina next-generation sequencing (NGS). **e.** DNA-level editing of target locus assessed by Sanger sequencing and ICE analysis. Editing efficiencies plotted here are the ‘ICE’d’ scores generated from ICE analysis. Non-treated samples were used as a control for analysis and are not displayed. **f.** Cell viability assessed by CellTiter-Glo assay one day after editing; reported in relative light units (RLU). X-axis labels indicate dose of editing reagents: 100 + 1 refers to 100 pmol RNP and 1 nmol peptide (10 μM peptide in 100 µL), 200 + 2 refers to 200 pmol RNP and 2 nmol peptide (20 μM peptide in 100 µL), and 400 + 4 refers to 400 pmol RNP and 4 nmol peptide (40 μM peptide in 100 µL). NT is non-treated. %NHEJ is non-homologous end-joining. *n*=3 biological blood donors. Error bars represent S.D. (standard deviation).

**Figure 3.**
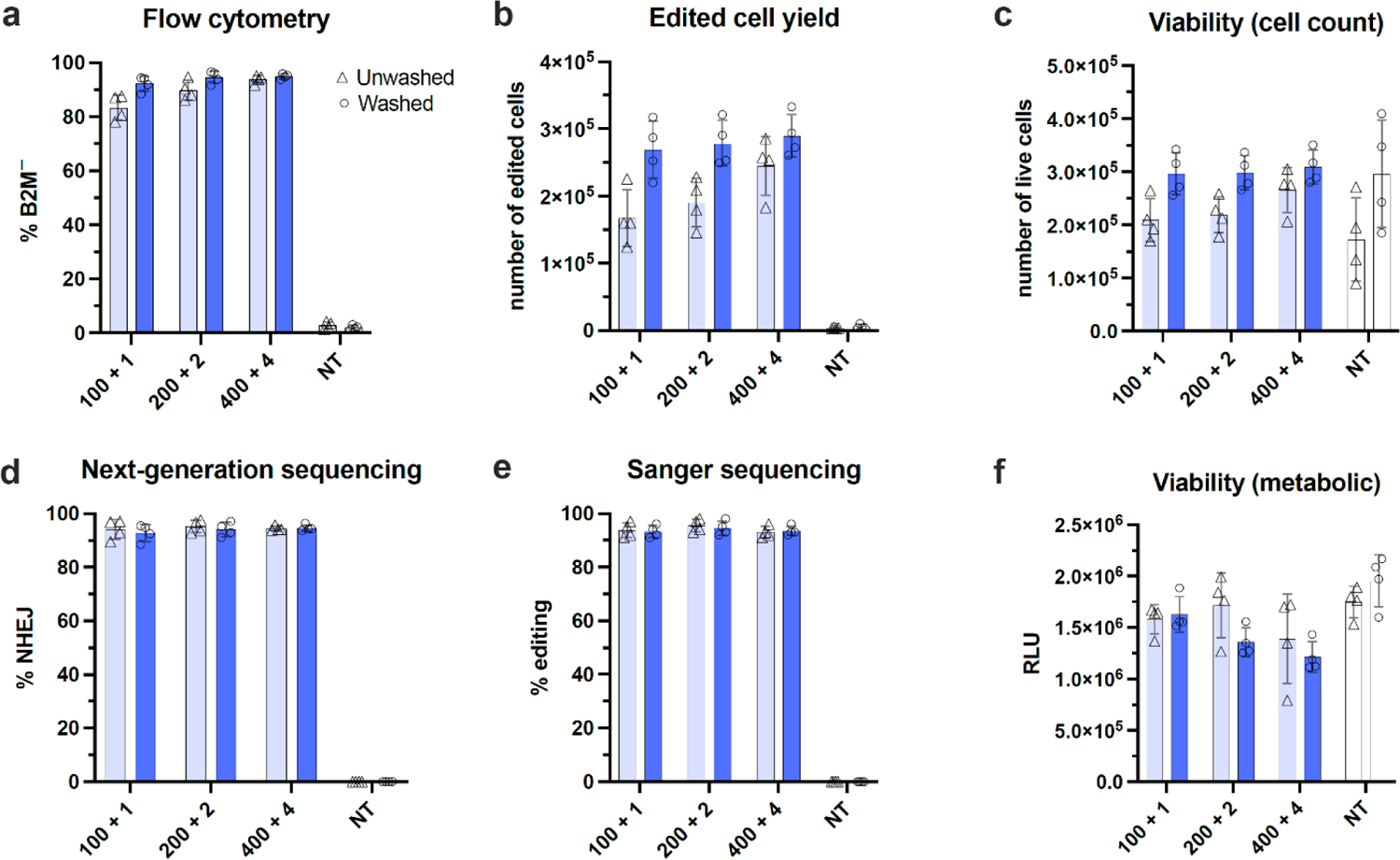
Cas12a-mediated editing of primary human T cells using PERC. Primary human T cells were incubated with editing reagents (Cas12a-Ultra-2xNLS) for 1 hour in Opti-MEM followed by either a 4x dilution (unwashed treatment) or washing off editing reagents (washed treatment) and recovering cells in growth medium supplemented with 5% FBS and 300U/mL IL-2. **a.** Protein-level KO of B2M assayed using flow cytometry. **b.** Number of cells showing a knockout in B2M surface expression, as assayed in (a). **c.** Cell viability depicted as number of live cells stained by Ghost Dye Red 780 four days after editing. **d.** Gene level editing of target locus assessed by Illumina next-generation sequencing (NGS). **e.** Gene level editing of target locus assessed by Sanger sequencing followed by ICE analysis. Editing rates plotted here are the ‘ICE’d’ scores generated from ICE analysis. Untreated samples were used as a control to run TIDE analysis, hence not displayed on the plot. **f.** Cell viability assessed by CellTiter-Glo assay two days after editing and reported in relative light units (RLU). X-axis labels indicate dose of editing reagents: 100 + 1 refers to 100 pmol RNP and 1 nmol peptide (10 μM peptide in 100 µL), 200 + 2 refers to 200 pmol RNP and 2 nmol peptide (20 μM peptide in 100 µL), and 400 + 4 refers to 400 pmol RNP and 4 nmol peptide (40 μM peptide in 100 µL). NT is non-treated. %NHEJ is non-homologous end-joining. *n*=4 biological blood donors. Error bars represent S.D. (standard deviation).

**Figure 4.**
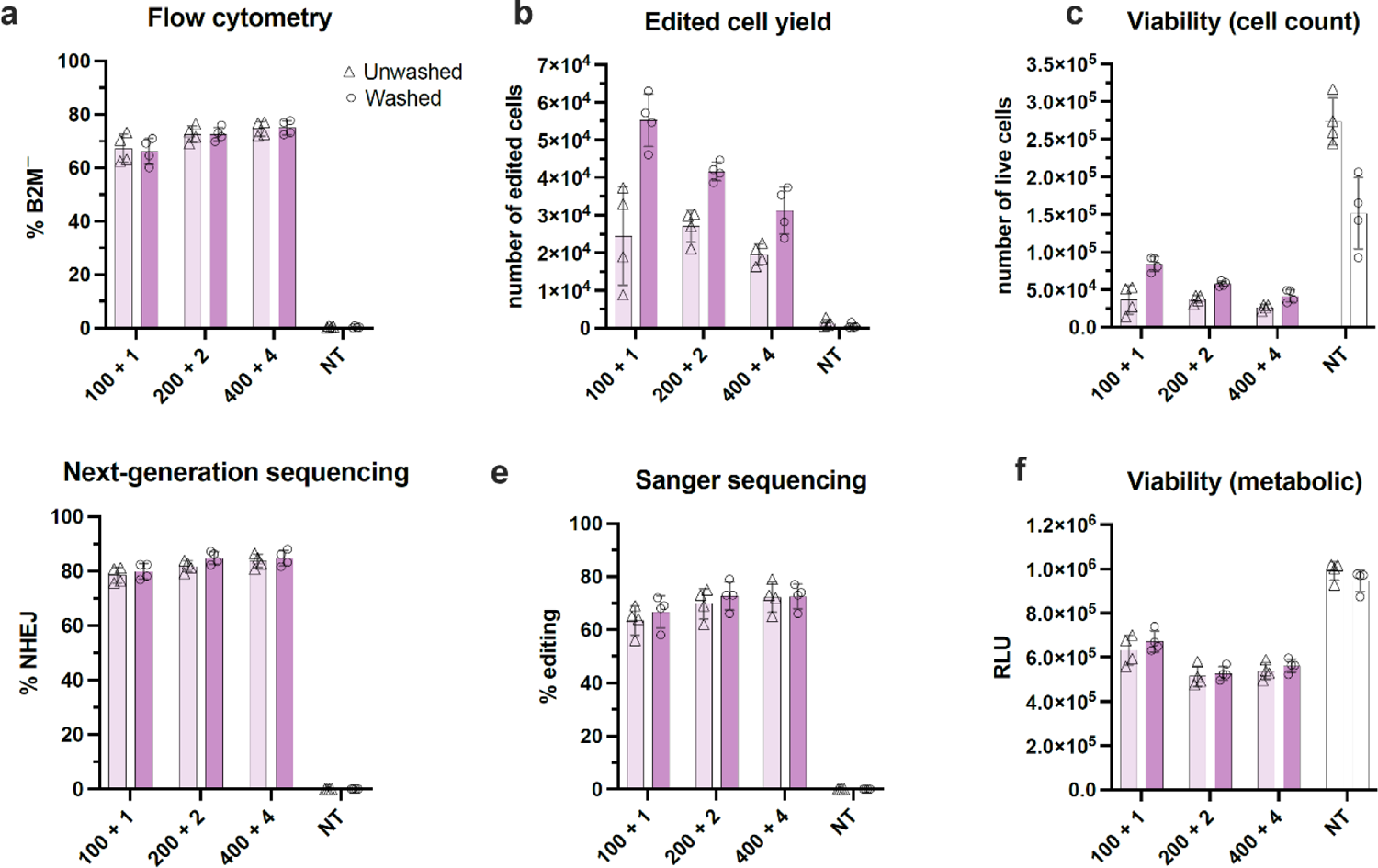
Cas9-mediated editing of primary HSPCs using PERC. HSPCs were incubated with editing reagents for 1 hour followed by either a 4x dilution in fresh HSPC culture medium (unwashed treatment) or washing off editing reagents (washed treatment) and recovering cells in fresh HSPC culture medium. **a.** Protein level knockout (KO) of target locus beta-2-macroglobulin (B2M) assayed using flow cytometry. **b.** Number of cells showing a knockout in B2M surface expression, as assayed in (a). **c.** Cell viability depicted as number of live cells stained by 7-AAD four days after editing. **d.** Gene level editing of target locus assessed by Illumina next-generation sequencing (NGS). **e.** Gene level editing of target locus assessed by Sanger sequencing followed by ICE analysis. Editing rates plotted here are the ‘ICE’d’ scores generated from ICE analysis. Untreated samples were used as a control to run TIDE analysis, hence not displayed on the plot. **f.** Cell viability assessed by CellTiter-Glo assay one day after editing and reported in relative light units (RLU). X-axis labels indicate dose of editing reagents: 100 + 1 refers to 100 pmol RNP and 1 nmol peptide (10 μM peptide in 100 µL), 200 + 2 refers to 200 pmol RNP and 2 nmol peptide (20 μM peptide in 100 µL), and 400 + 4 refers to 400 pmol RNP and 4 nmol peptide (40 μM peptide in 100 µL). NT is non-treated. %NHEJ is non-homologous end-joining. *n*=4 biological blood donors. Error bars represent S.D. (standard deviation).

**Figure 5.**
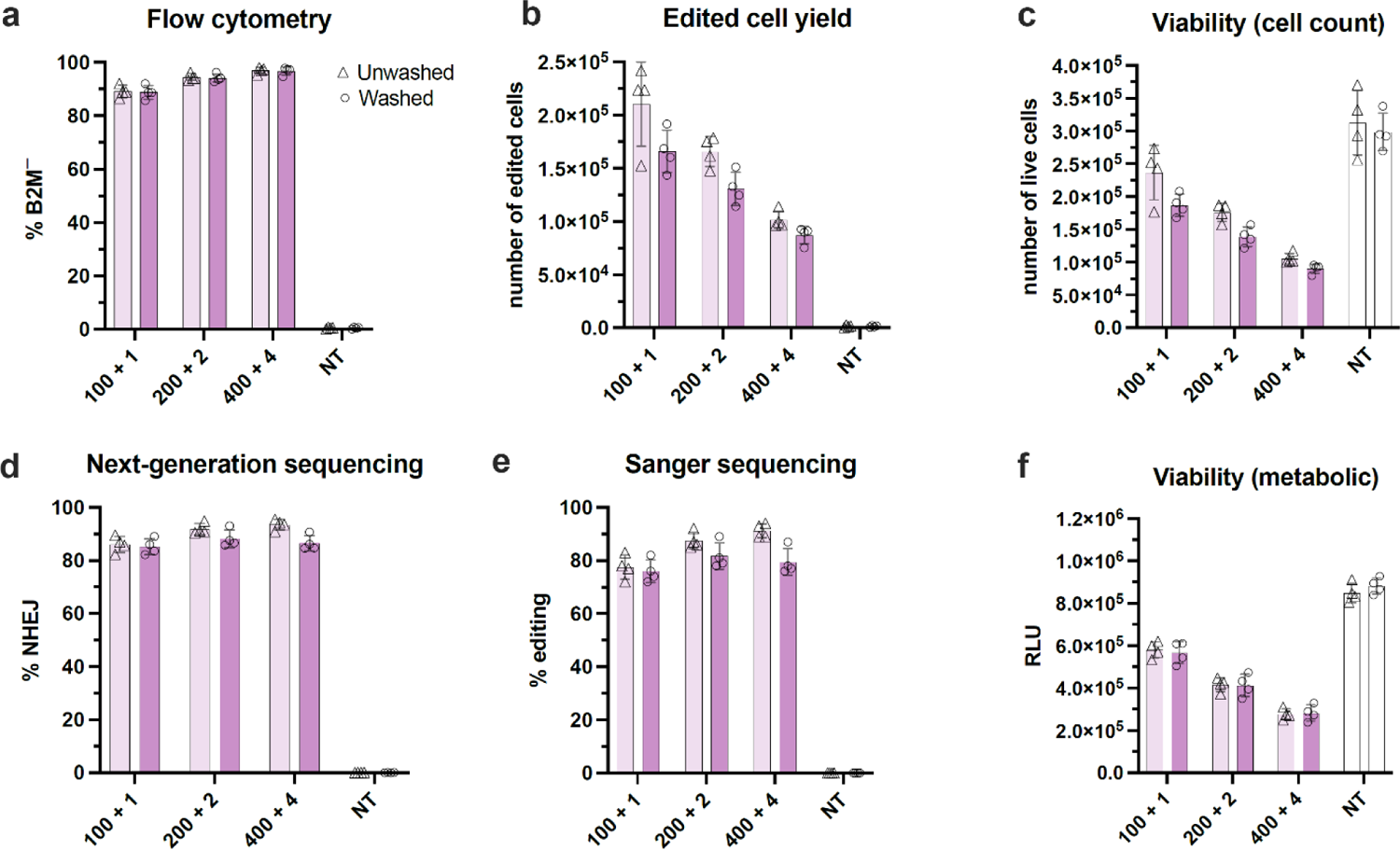
Cas12a-mediated editing of primary human HSPCs using PERC. HSPCs were incubated with editing reagents for 1 hour followed by either a 4x dilution in fresh HSPC culture medium (unwashed treatment) or washing off editing reagents (washed treatment) and recovering cells in fresh HSPC culture medium. **a.** Protein level knockout (KO) of target locus beta-2-macroglobulin (B2M) assayed using flow cytometry. **b.** Number of cells showing a knockout in B2M surface expression, as assayed in (a). **c.** Cell viability depicted as number of live cells stained by 7-AAD four days after editing. **d.** Gene level editing of target locus assessed by Illumina based next-generation sequencing (NGS). **e.** Gene level editing of target locus assessed by Sanger sequencing followed by ICE analysis. Editing rates plotted here are the ‘ICE’d’ scores generated from ICE analysis. Untreated samples were used as a control to run TIDE analysis, hence not displayed on the plot. **f.** Cell viability assessed by CellTiter-Glo assay one day after editing and reported in relative light units (RLU). X-axis labels indicate dose of editing reagents: 100 + 1 refers to 100 pmol RNP and 1 nmol peptide (10 μM peptide in 100 µL), 200 + 2 refers to 200 pmol RNP and 2 nmol peptide (20 μM peptide in 100 µL), and 400 + 4 refers to 400 pmol RNP and 4 nmol peptide (40 μM peptide in 100 µL). NT is non-treated. %NHEJ is non-homologous end-joining. *n*=4 biological blood donors. Error bars represent S.D. (standard deviation).

**Figure 6.**
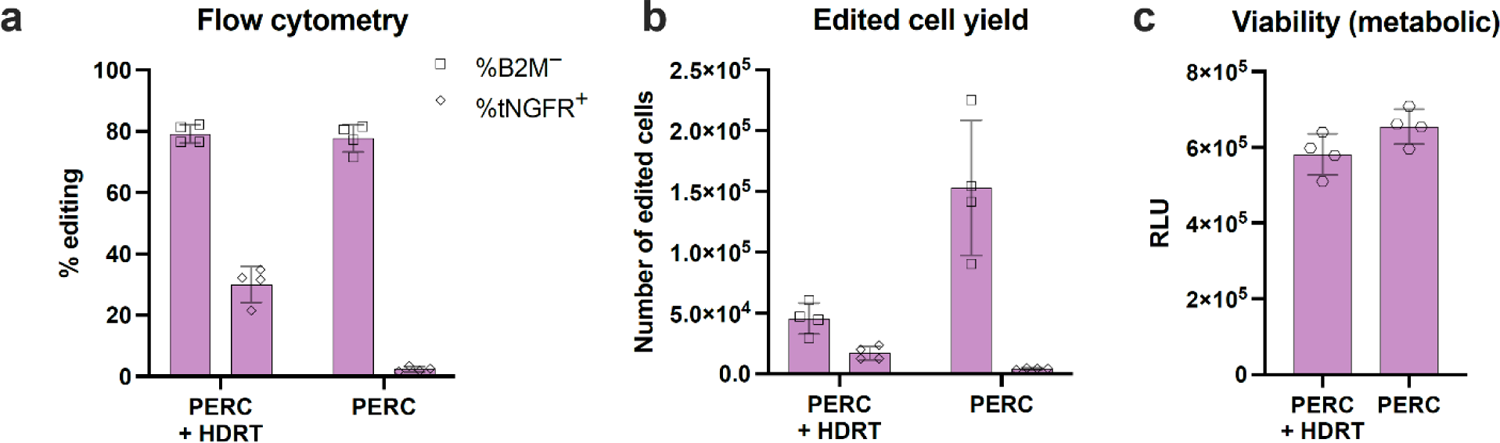
Cas12a-mediated knock-in via HDR in primary human HSPCs using PERC. HSPCs were incubated with 100 pmol Cas12a RNP and 10 μM peptide for 1 hour followed by washing off editing reagents and recovering cells in fresh HSPC culture medium. AAV6 packaged with HDR template (HDRT) was added to the recovered cells, which were incubated overnight. **a.** Protein-level KO of B2M paired with (PERC+HDRT) or without (PERC) knock-in (KI) of tNGFR assayed using flow cytometry. In the HDRT condition, %B2M^−^ does not exclude cells that are also tNGFR^+^. **b.** Number of cells bearing the intended edits using PERC + HDRT or PERC alone, as assayed in (a). **c.** Cell viability assessed by CellTiter-Glo assay one day after editing and reported in relative light units (RLU). *n*=4 biological blood donors. Error bars represent S.D. (standard deviation).

## MATERIALS & METHODS

### Biological material

For sourcing human T cells, pre-isolated cells can be purchased, or cells can be isolated from peripheral blood leukopaks.

- Human peripheral blood pan T cells (STEMCELL Technologies #70024)
- Human peripheral blood leukopak (STEMCELL Technologies #70500.1), half-size leukopak, fresh)

Leukopaks need to be scheduled to arrive on a day when there is time to process the pack. We do not recommend using frozen leukopaks, as they can result in unhealthy cells and low yields after processing.

Frozen human CD34⁺ HSPCs isolated from mobilized peripheral blood can be purchased from multiple vendors (AllCells #mLP RegF CR CD34+ PS 1M), (STEMCELL Technologies #70060.1). We have used CD34⁺ HSPCs mobilized via G-CSF, Plerixafor, or G-CSF and plerixafor.

### !CAUTION

Human blood samples should be handled according to approved biosafety protocols and regulations set by each institution for protection of patient confidentiality and health information and user protection. Leukopak donors are typically tested for HIV and hepatitis A and B. Handling scientists are typically required to be trained for handling potential bloodborne pathogens and additionally recommended to be vaccinated against hepatitis A and B.

### Reagents

Isolating primary human T cells (optional):

- Dulbecco’s phosphate buffered saline (DPBS), no calcium, no magnesium (Fisher Scientific #14190144)
- Fetal bovine serum (FBS) (Thermo Fisher Scientific #26140079)
- Ultrapure 0.5 M ethylenediamine tetraacetic acid EDTA, pH 8.0 (Invitrogen #15575-038)
- Lymphoprep (STEMCELL Technologies #07861)
- Trypan blue stain 0.4% (Fisher Scientific # T10282)
- EasySep negative isolation kit for CD3^+^ cells (STEMCELL Technologies #19051)
- Bambanker (Wako chemicals, Fisher Scientific #NC9582225), CryoStor (BioLife solutions #CS10) or equivalent

Cell culture:

- Autoclaved water
- Aqua-Clear water conditioner (SP Scienceware #F17093-0000)
- Ethanol 200 proof (Koptec #V1001)

**!CAUTION** Ethanol is flammable. Store in secondary container in a dedicated cabinet for flammable reagents.

- Germicidal Ultra Bleach (Waxie 170018)

**!CAUTION** Bleach is a skin irritant and can cause skin burns or eye damage. Use proper protective equipment (PPE) when handling and store in a secondary container.

Culturing primary human T cells:

- X-VIVO 15 serum-free hematopoietic cell medium (Lonza #04-418Q)
- FBS (Thermo Fisher Scientific #26140079)
- 2-mercaptoethanol 1,000× 55 mM in DPBS (Gibco #21985-023)

**!CAUTION** Toxic if swallowed or inhaled. Use PPE when handling and store in a secondary container.

- N-acetyl L-cysteine (Millipore Sigma #A9165-25G)
- Recombinant human IL-2 (Peprotech #200-02)
- Recombinant human IL-7 (Peprotech #200-07)
- Recombinant human IL-15 (Peprotech #200-15)
- Bovine serum albumin solution (Sigma-Aldrich #A7979-50ML)
- Dynabeads™ human T-activator CD3/CD28 for T cell expansion and activation (Thermo Fisher Scientific # 11132D)

Culturing primary human hematopoietic stem cells:

- StemSpan™ SFEM II (STEMCELL Technologies # 09655)
- StemSpan™ CC110 (premixed cytokine cocktail) (STEMCELL Technologies #02697); or, recombinant human TPO (PeproTech #300-18), recombinant human Flt3-ligand (PeproTech #300-19), and recombinant human SCF (PeproTech #300-07)
- Bovine serum albumin solution (for cytokine preparation; Sigma-Aldrich #A7979-50ML)
- Penicillin/streptomycin (Pen-Strep) (optional; Fisher Scientific #15140-122)

Assembly of CRISPR RNP:

- ELIMINase decontaminant (Fisher Scientific #04-355-2)
- DEPC-treated water (Growcells #UPW-0125)
- 1 M HEPES buffer solution (Gibco #15630-080)
- 5 M sodium chloride (Invitrogen #AM9760G)
- 1 M magnesium chloride (Fisher Scientific #BP214-500)
- 1 M D-(+)-Trehalose, dihydrate (Fisher Scientific #T0331-500G)
- gRNA or crRNA (self-synthesized, or synthesized by IDT or Synthego)
- CRISPR protein (purified in-house, or procured from commercial vendors or UC Berkeley QB3 MacroLab)

- Cas9 (choose from the following options):

- Cas9-triNLS (contact UC Berkeley MacroLab at https://qb3.berkeley.edu/facility/qb3-macrolab/)
- TrueCut^TM^ Cas9 protein v2 (Invitrogen #A36496)
- Cas9-6xNLS (Addgene # 88917 and #196246)
- Alt-R® S.p. Cas9 nuclease V3 (IDT # 1081058)
- Alt-R® S.p. HiFi Cas9 nuclease V3 (IDT #1081061)
- SpCas9 2xNLS nuclease (Synthego)
- Cas12a (choose from the following options):

- Alt-R® A.s. Cas12a (Cpf1) Ultra (IDT #10007804)
- A.s.Cas12a-Ultra-2xNLS (email UC Berkeley MacroLab at)
- A.s.Cas12a-Ultra-5xNLS (Addgene #218775)

**CRITICAL** Proteins should be flash-frozen and stored at −80°C. Aliquot for convenient use and avoid multiple freeze-thaws. For more information on storage conditions, refer to **Box 1**.

#### Box 1

##### Storage and stability of editing reagents

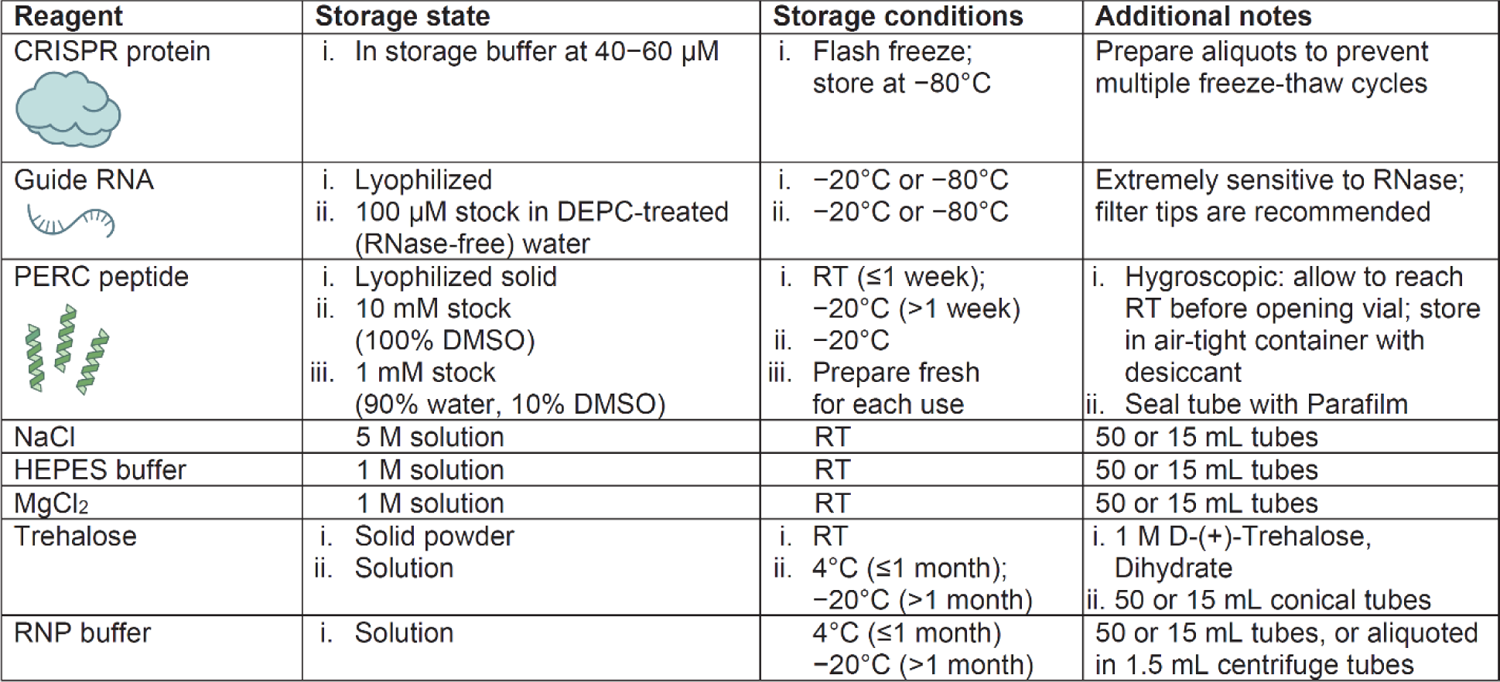

Preparing peptide:

- DMSO (Fisher Bioreagents #BP231-100)

**!CAUTION** flammable and toxic, use gloves to handle

- Peptide INF7TAT-P55 (CPC Scientific 1 mg #IMMO-008A, 5 mg #IMMO-008B, or custom synthesis)
- Peptide INF7TAT-A5K (CPC Scientific 1 mg #IMMO-007A, 5 mg #IMMO-007B, or custom synthesis)

**!CAUTION** use gloves when handling these peptides

Coincubation of cells and editing reagents (for T cells):

- Opti-MEM reduced serum medium (Thermo Fisher Scientific #11058021)

Analysis of editing outcomes:

- CellTiter-Glo^®^ luminescent cell viability assay (Promega #G9242)
- Quick Extract buffer (Biosearch Technologies #QE09050)
- Primestar GXL polymerase (Takara #R050A)
- AMPure XP beads (Beckman Coulter #A63880) or equivalent SPRI beads
- DEPC-treated water (Growcells #UPW-0125)
- Ethanol 200 proof (Koptec #V1001)

**!CAUTION** Ethanol is flammable. Store in secondary container in a dedicated cabinet for flammable reagents.

- MiSeq reagent kit v2 (300 cycles) (Illumina #MS-103-1002)
- Agarose ultrapure (Fisher Scientific #16500-100)
- SYBR Safe DNA gel stain (Fisher Scientific #S33102)
- Tris-acetate-EDTA (TAE) Buffer, 10X (Fisher Scientific #BP13351)
- Ghost Dye Red 780 (Tonbo Biosciences #13-0865-T500) or 7-AAD staining solution (Fisher Scientific # 559925)

Antibodies for staining of T cells for flow cytometry analysis:

- Anti-human CD3-PE (BioLegend #300408, RRID:AB_314062 Clone UCHT1)
- Anti-human CD4-Brilliant Violet 421™ (BioLegend #317434, RRID:AB_2562134 Clone OKT4)
- Anti-human CD8a-APC (BioLegend #300912, RRID:AB_314116 Clone HIT8a)
- Anti-human beta-2-microglobulin-FITC (BioLegend #316304, RRID:AB_492837 Clone 2M2)

Antibodies for immunophenotyping HSPCs for flow cytometry analysis:

- Anti-human CD34-Brilliant Violet 421™ (BioLegend #343610, RRID:AB_2561358 Clone 561)
- Anti-human beta-2-microglobulin-APC (BioLegend #316312, RRID:AB_10641281 Clone 2M2)
- Anti-human beta-2-microglobulin-FITC (BioLegend #316304, RRID:AB_492837 Clone 2M2)
- Anti-human NGFR-FITC (BioLegend #345104, RRID:AB_2282828 Clone ME20.4)

### Equipment

Cell culture:

- Biosafety Level 2 (BSL-2) laminar flow hood (Thermo Fisher scientific #1300 Series A2 or equivalent)
- Cell culture incubator (Thermo Fisher Scientific HERAcell vios160i or equivalent)
- Water bath (Fisherbrand Isotemp GPD05 or equivalent)
- Mini centrifuge (Genesee Scientific #31-501 or equivalent)
- Benchtop centrifuge (Beckman Allegra X-12R or equivalent)
- Microplate carrier for benchtop centrifuge (VWR # BK392806)
- Optical light microscope
- T75 and T25 tissue culture treated filter cap flasks (USA Scientific #CC7682-4875, #CC7682-4825 or equivalent)
- 50 mL conical tubes (Fisher Scientific #352070 or equivalent)
- 15 mL conical tubes (Fisher Scientific # 352196 or equivalent)
- Standard microcentrifuge tubes (Thomas Scientific #1149K01 or equivalent)
- Reagent reservoirs (Fisher Scientific #3054-1006, 3054-2012 or equivalent)
- Polystyrene serological pipettes 5, 10, 25, 50 mL (Fisher Scientific #4487, 4488, 4489, 4490, or equivalent)
- Pipette L1000, L200, L20, L10, L2 (Rainin #17014382, 17013805, 17014388, 17014392, 17014393 or equivalent)
- Multi-channel pipette (Rainin #17013810, 17013808, or equivalent)
- Filter tips (Rainin #3038912,17005859, 30389225, or equivalent)
- Pipet gun/filler (Thermo Fisher Scientific #9511 or equivalent)
- Stripettes (disposable polystyrene serological pipettes) 50 mL, 25 mL, 10 mL
- T150 flask, T75, T25 (USA Scientific #CC7682-4815, #CC7682-4875, #CC7682-4825)
- Hemocytometer counting chamber (Fisher Scientific #02-671-51B)
- Liquid nitrogen tank (Thermo Fisher Scientific, Locator™ Plus Rack and Box Systems #CY509109 or similar)

Isolating primary human T cells:

- Argos Technologies aspirating serological pipette volume 10 mL (Fisher Scientific #3395167)
- Easy50 EasySep^TM^ magnet (STEMCELL Technologies #18002)
- Cryogenic vials, internal thread (Corning 1.2 mL #CLS430487)
- Mr. Frosty, Corning CoolCell cell freezing container (#432138) or equivalent
- Freezer −80°C

Culturing primary human T cells:

- Stericup vacuum filtration system (Millipore #S2GPU05RE)
- Dynamag^TM^-15 magnetic stand (Thermo Fisher Scientific #12301D)
- Corning Falcon 96-well round-bottom tissue culture-treated surface cell culture plate (Millipore Sigma #CLS353077)

Culturing primary human HSPCs:

- Steriflip-GP 0.22 µm (Millipore #SCGP00525 or #S2GPU05RE)
- 96-well flat-bottom tissue-culture treated plates (Corning Costar #3599))

Assembly of CRISPR RNP:

- Standard microcentrifuge tubes (Thomas Scientific #1149K01 or equivalent)
- Mini dry bath (Fisher scientific #14-955-221)
- Vortex mixer (Fisher Scientific #02-215-414 or similar)
- Mini centrifuge (Genesee Scientific #31-501 or similar)
- Syringe 10 mL (Fisher Scientific #14-955-459 or similar)
- 0.2 µm sterile syringe filter (VWR #28145-501 or similar)
- Amicon Ultra centrifugal filter device (Millipore-Sigma #UFC505024) Preparing peptide:
- Standard microcentrifuge tubes (Thomas Scientific #1149K01 or equivalent)
- 50 mL conical tubes (Fisher Scientific #352070 or equivalent)
- Kimwipes^TM^ (VWR #21903-005 or equivalent)
- Benchtop Centrifuge (Beckman Allegra X-12R or equivalent)
- Airtight container or jar (Thermo Fisher Scientific #2118-0032 or similar)
- Drierite, indicating desiccant (Fisher Scientific #075783A)
- Parafilm (VWR #52858-076)

Analysis of editing outcomes:

- Thermocycler (Bio-Rad #T100 or equivalent)
- SPRI ring super magnet plate (Beckman Coulter #A32782)
- Polypropylene 0.2 mL 96-well PCR plates (USA Scientific #1402-9598)
- Hard-shell 96-well PCR plates (Bio-Rad #HSS9601)
- Plate sealing foil (USA Scientific #2923-0110)
- NanoDrop One (Thermo Fisher Scientific)
- Gel electrophoresis unit (Bio-Rad #1704487)
- Power unit (Bio-Rad #1645050)
- Gel casting tray (Bio-Rad #1704436)
- Plate reader (Tecan Spark or equivalent)
- Flow cytometer (Attune NxT, or equivalent)
- FlowJo software license (optional)
- Sequencing partner (Quintara or equivalent)
- CRISPResso (http://crispresso2.pinellolab.org/submission)
- ICE (https://ice.synthego.com)
- TIDE (https://tide.nki.nl/)

### Reagent setup

- EasySep^TM^ buffer

Prepare EasySep buffer for use during leukopak isolation of T cells by combining the following ingredients in a clean BSL-2 hood:

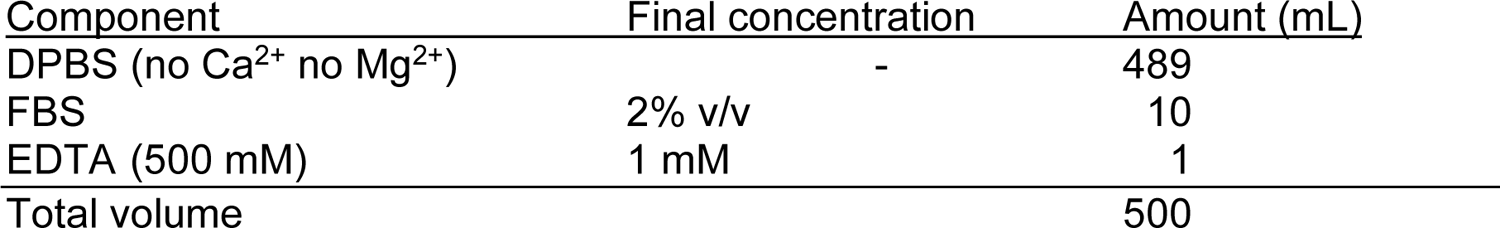

 EasySep buffer can be stored at 4°C for up to a month.

- T cell growth medium

Prepare growth medium for T cell culture by combining X-VIVO 15 medium (Lonza), 5% fetal bovine serum, 50 μM β-mercaptoethanol, and 10 mM N-acetyl-L-cysteine in a clean BSL-2 hood. Follow the table below. Sterile-filter using a Stericup bottle top filter.

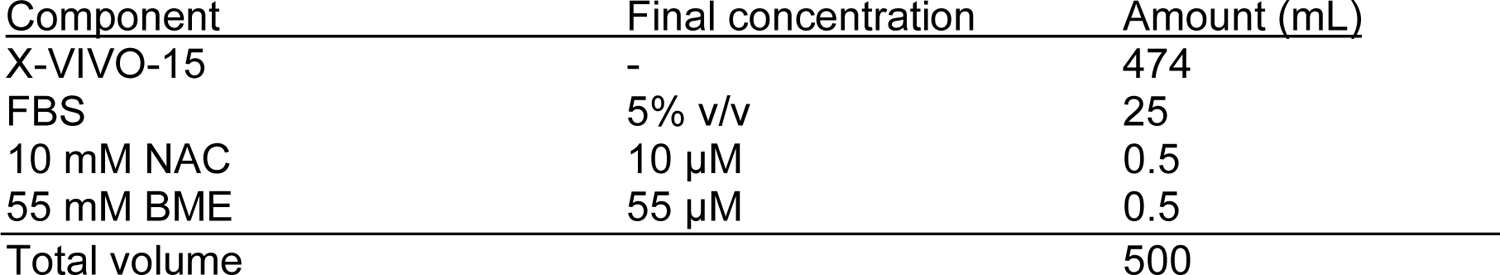

**CRITICAL** Prepared medium can be stored at 4°C for up to a month.

- Bovine serum albumin solution: prepare 10 mL of 0.1% w/v solution in 1× DPBS and sterile-filter using a 0.22 µm filter. This solution is used to resuspend cytokines. We recommend preparing this fresh, as it will be used to make cytokine stocks for long-term storage.
- Human IL-2 stocks: lyophilized cytokine can be resuspended to make 500,000 U/mL stock in 0.1% w/v bovine serum albumin solution in 1× DPBS. Sterile-filter, aliquot, and store at 4°C for up to four weeks or −80°C long-term.
- Human IL-7 stocks: lyophilized cytokine can be resuspended to make 5 µg/mL stock in 0.1% w/v bovine serum albumin solution in 1× DPBS. Sterile-filter, aliquot, and store at 4°C for up to four weeks or −80°C long-term.
- Human IL-15 stocks: lyophilized cytokine can be resuspended to make 5 µg/mL stock in 0.1% w/v bovine serum albumin solution in 1× DPBS. Sterile-filter, aliquot, and store at 4°C for up to four weeks or −80°C long-term.
- HSPC culture medium

Combine StemSpan^TM^ SFEM II with SCF (Stem-cell factor), TPO (Thrombopoietin) and Flt3L (Fms-related tyrosine kinase 3 ligand). Adding Pen-Strep is optional. Scale according to the volume needed.

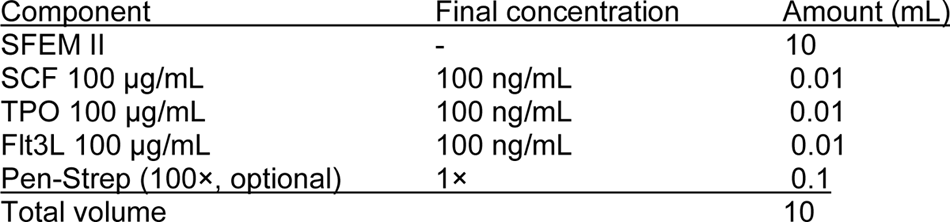

Alternatively, StemSpan CC110 can be used, which is a premixed cytokine cocktail of the individual cytokines listed above.

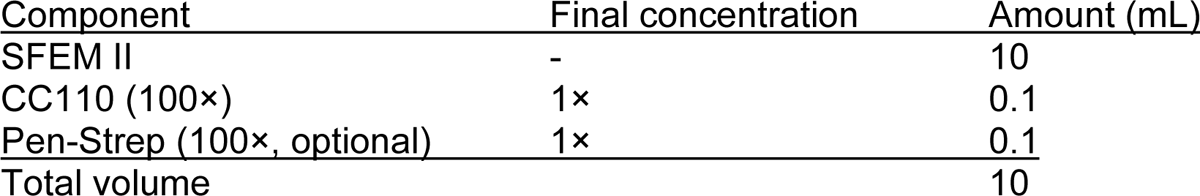

**CRITICAL** Prepared HSPC culture medium can be stored at 4°C for up to a week. StemSpan^TM^ SFEM II comes as 100-mL or 500-mL bottles and can be stored at −20°C for up to 18 months from the date of manufacture or at 2–8°C for up to a month. Refreezing is not recommended. Aliquot and freeze the medium in smaller batches so that the appropriate volume can be thawed and prepared when needed. Mix thoroughly before aliquoting. If the full bottle is to be stored at 4°C for a month, sterile-filter the medium obtained from the bottle before preparing the HSPC culture medium. Prepare appropriately sized aliquots of StemSpan CC110 if the full 1 mL is not planned to be used within 1 month at 4°C. Store the aliquots at −20°C.

- SCF, TPO, and Flt3L cytokines are resuspended to make 100 µg/mL stock in 0.1% Bovine serum albumin solution in 1×DPBS. Aliquot and store at 4°C for up to 2 weeks, or at −20°C or −80°C long-term.
- Prepare appropriately sized aliquots of Pen-Strep. Store at 4°C for up to 1 month, or −20°C long-term.
- RNP buffer:

Prepare buffer to assemble RNP in by combining the reagents below. Sterile-filter using a 0.22 µm syringe filter. The buffer can be stored at 4°C for up to 4 weeks or stored frozen at −20°C for several months.

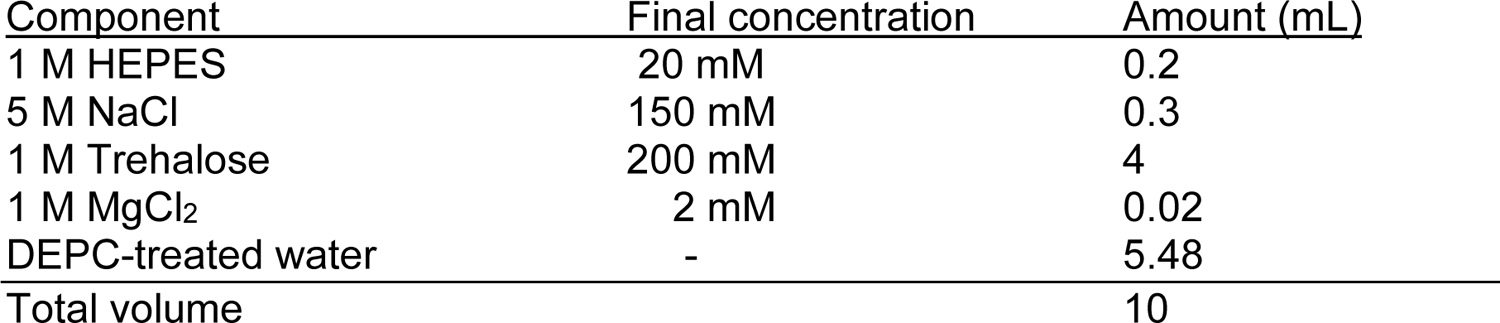

The addition of MgCl2 is optional and can improve editing efficiency in certain cell types. The concentration of trehalose should be 100 mM in final RNP volume. The above concentration of trehalose is assuming the Cas protein is at 40 µM. If Cas protein is at a different concentration, first adjust Cas to 40 µM using base buffer (with 20 mM HEPES and 150 mM NaCl in water) or calculate amount of trehalose to add to ensure a final concentration of 100 mM.

**CRITICAL** The pH of all the solutions should be around 7.0, which can be quickly tested using pH indicator strips. We often find the prepared buffer to be already at pH 7, avoiding the need for any further adjustment.

gRNA resuspension

- Bring a vial of lyophilized gRNA to room temperature.
- Briefly spin down the tube before opening to ensure all lyophilized guide is at the bottom of the tube.
- To make a 100 µM stock of gRNA, resuspend 10 nmol of RNA in 100 µL of DEPC-treated nuclease-free water (scale as needed).
- Pulse-vortex for 30 seconds after adding water to the lyophilized RNA and pipette to mix well. Briefly spin down vial one more time.
- The gRNA stock can be used immediately or stored at −20°C for several months, or −80°C for longer. Aliquot into small volumes to avoid multiple freeze-thaws. For more information, refer to **Box 1**.

Preparing peptide stock

- Peptides purchased from CPC Scientific are typically in lyophilized form. Peptides should be stored wrapped in parafilm in an airtight container with desiccant at −20°C. Refer to **Box 1** for more on storage recommendation.
- Before opening, leave the lyophilized vial at room temperature for 20 minutes. Spin down the vial wrapped in a Kimwipe and inside of a 50 mL conical tube at 2000 × *g* for 1 minute.
- Carefully break the seal of the vial and resuspend the peptide in 100% DMSO to prepare a stock concentration of 10 mM. Calculate the volume of DMSO to use based on the molecular weight of the peptide:
- It is important to be careful when opening and handling the vial of lyophilized peptide to avoid the powder from blowing out. Be careful not to touch the inside of the lid. Replace the lid securely and roll or invert the vial to make sure any lyophilized material in the neck or lid is also resuspended.
- Centrifuge the resuspended vial one more time. Aliquot into small working volumes and store at −20°C.

**CRITICAL** Peptide stock should be stored at −20°C in a desiccator, tightly parafilm-sealed, and kept away from moisture to maintain long-term potency. If a desiccator is not available, we recommend using an air-tight jar and sealing the tube with parafilm. Peptide is stable in 100% DMSO for several months.

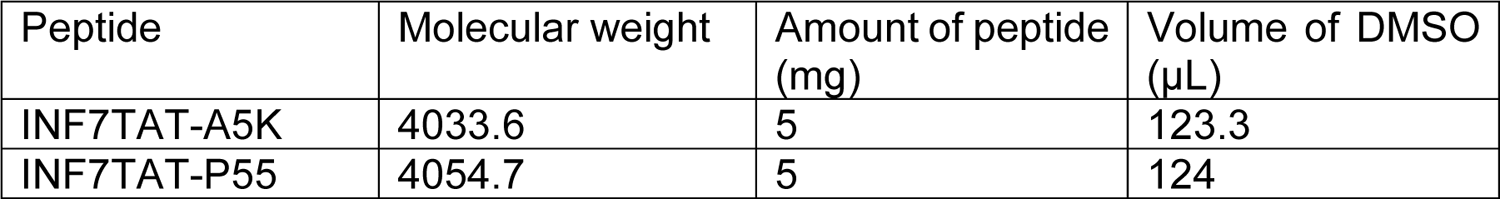

Coincubation of cells with editing reagents:

- Recovery medium (1.5×, for T cells)

**CRITICAL** Cytokines and FBS should be added fresh to X-VIVO15 medium the same day.

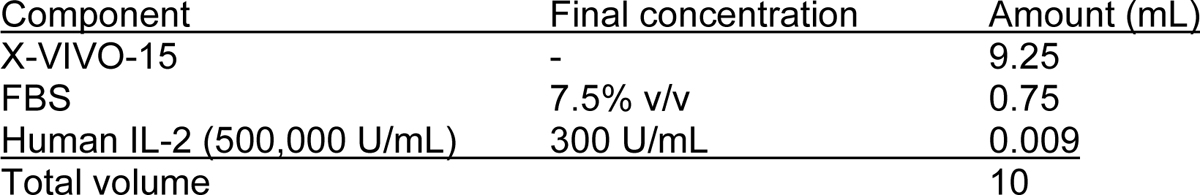

Assessing editing efficiency:

- Flow cytometry buffer

Make 500 mL of buffer to use for washing and preparing cells for flow cytometry by supplementing 1× DPBS (no calcium, no magnesium) with 2% fetal bovine serum and EDTA using the table below. This buffer can be stored at 4°C for up to 4 weeks.

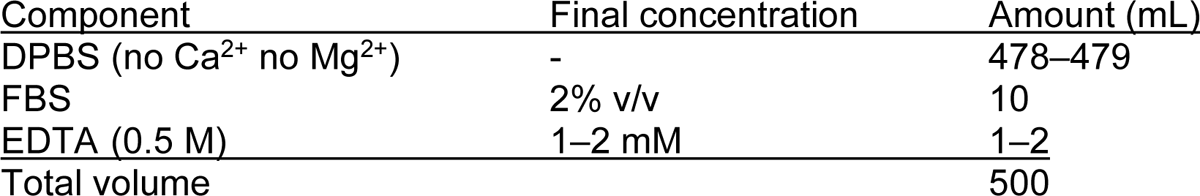

## PROCEDURE

### Culturing cells of interest

1. For culturing and preparing cells for editing: if working with T cells, follow option (A), and if working with HSPCs, follow option (B).

A. Isolating and culturing primary human T cells

Isolation of primary human T cells from leukapheresis packs Timing 5–6 hours

i. Let reagents reach room temperature before starting.
ii. Follow manufacturer’s recommended protocol to isolate peripheral blood mononuclear cells (PBMCs)^82^.
iii. Take a 10 µL sample to count cells. Pipette 10 µL of trypan blue solution onto the cells, mix well, and load a hemocytometer counting chamber with 10 µL of the 1:1 diluted sample. Count four corners of the grid and average the count. Calculate the concentration of the cells using the formula below: Average cell count × dilution factor × 10,000 = cell concentration (cells/mL) **CRITICAL STEP** Automated counting machines are not very accurate for PBMCs, especially for naive, non-stimulated cells. We recommend always counting PBMCs or T cells manually using a hemocytometer.
iv. Isolate bulk T cells from PBMCs by magnetic negative selection using EasySep isolation kits for human T cells (STEMCELL) per the manufacturer-provided instructions.
v. Count the number of cells using a hemocytometer to determine the yield of T cells obtained. Take a 10 µL sample, pipette equal volume of trypan blue solution onto the cells, mix well, and load a hemocytometer counting chamber with 10 µL of the 1:1 diluted sample. Count four corners of the grid and average the count. Calculate the concentration of the cells using the formula below: [average cell count] × [dilution factor] × 10,000 = cell concentration (cells/mL)
vi. Isolated cells can be directly activated for use in an experiment approximately 48 hours later. **Pause point** Alternatively, T cells can be stored in Bambanker or CryoStor freezing medium (or equivalent) in cryovials stored in a liquid nitrogen tank for use later. Culturing and preparing primary human T cells for editing Timing 2–3 hours hands on, 3 days of culture *Day −3 (relative to the day of editing)* **Timing 30 minutes** Frozen T cells should be thawed three days before the day of the editing experiment (day −3 relative to delivery) (**Fig. 1c**). It is recommended to culture T cells at a concentration of 1 × 10^6^ cells/mL. However, frozen cells should be allowed to rest overnight at a concentration of 1.5 × 10^6^ cells/mL to compensate for cells lost during the freeze-thaw process. For 50 × 10^6^ cells, culture in a volume of 33.3 mL of growth medium on the rest day. **CRITICAL STEP** Because biological donors often have varying activation rates and responses, we recommend conducting these experiments with at least two biological replicates.
vii. Prepare growth medium according to instructions provided under ‘reagent setup’.
viii. Pre-warm growth medium in a 37°C water bath.
ix. In a clean BSL-2 cabinet, prepare a 15 mL conical tube with 8 mL of growth medium.
x. Label a T25 flask with the date, your initials, and donor identification. Pipette 23.3 mL of prewarmed growth medium into the flask.
xi. Remove a cryovial of frozen T cells from liquid nitrogen storage and immediately place in the 37°C water bath. If the vial needs to be transported, do so on dry ice.
xii. Carefully monitor thawing of the frozen cell stock, removing it from the water bath when 90% thawed. This step takes approximately 3 minutes.
xiii. When cells have thawed completely, inside the biosafety cabinet, carefully unscrew the cap without touching the inside of the vial.
xiv. Pipette cells into growth medium previously prepared in the 15 mL conical tube.
xv. Wash the inside of the empty cryovial with 1 mL of growth medium to collect any remaining cells and add to the 15 mL tube.
xvi. Centrifuge at 300 × *g* for 5 minutes, RT to pellet the cells.
xvii. Pour off the supernatant and resuspend in 10 mL of warm growth medium.
xviii. Repeat the previous two steps to wash for a total of two times.
xix. Mix well by pipetting, and dispense cells into the prepared T25 flask.
xx. Place the T25 flask containing T cells in a tissue culture incubator at 37°C and 5% CO2. Let the cells rest overnight. **Pause point** Cells can be left in the incubator overnight until activation. *Day −2: Activation and expansion of T cells* **Timing 1–1.5 hours** Cells should be activated after ∼24 hours of rest or directly after isolation of T cells from a leukopak (day −2 relative to delivery).
xxi. Transfer cells into a 50 mL conical tube. Mix well using a 10 mL disposable serological pipette before transferring cells. Measure and take note of the volume of cells transferred. **CRITICAL STEP** The measured volume might be slightly less than the volume of the culture the previous day due to evaporation loss. We typically observe ∼1 mL loss.
xxii. Take a 10 µL sample to count cells as described previously.
xxiii. Centrifuge cells in the 50 mL tube at 300 × *g* for 5 minutes RT.
xxiv. Adjust the cell concentration to 1 × 10^6^ cells/mL, keeping the cells in the same medium from the previous day.
xxv. Cells might be at a concentration lower than 1 × 10^6^ cells/mL and some volume of medium will need to be removed. To adjust the concentration, centrifuge the cells at 300 × *g* for 5 minutes RT and carefully pipet out some volume of medium from the top surface without disturbing the cell pellet.
xxvi. If the cells are at a higher concentration and need to be diluted, add prewarmed growth medium on top.
xxvii. When centrifuging the cells, take a 1 mL medium aliquot from the supernatant, and set aside. **CRITICAL STEP** Do not dispose medium from the previous day. T cells secrete cytokines that aid in their growth and expansion, so it is important to reuse medium from the previous day.
xxviii. To activate T cells, culture at 1 × 10^6^ cells/mL for two days in supplemented X-VIVO 15 medium with anti-human CD3/CD28 magnetic Dynabeads at a bead-to-cell ratio of 1:1, 200 U/mL, IL-2, 5 ng/mL IL-7, and 5 ng/mL IL-15. The table below shows an example calculation for activating 30 × 10^6^ cells:

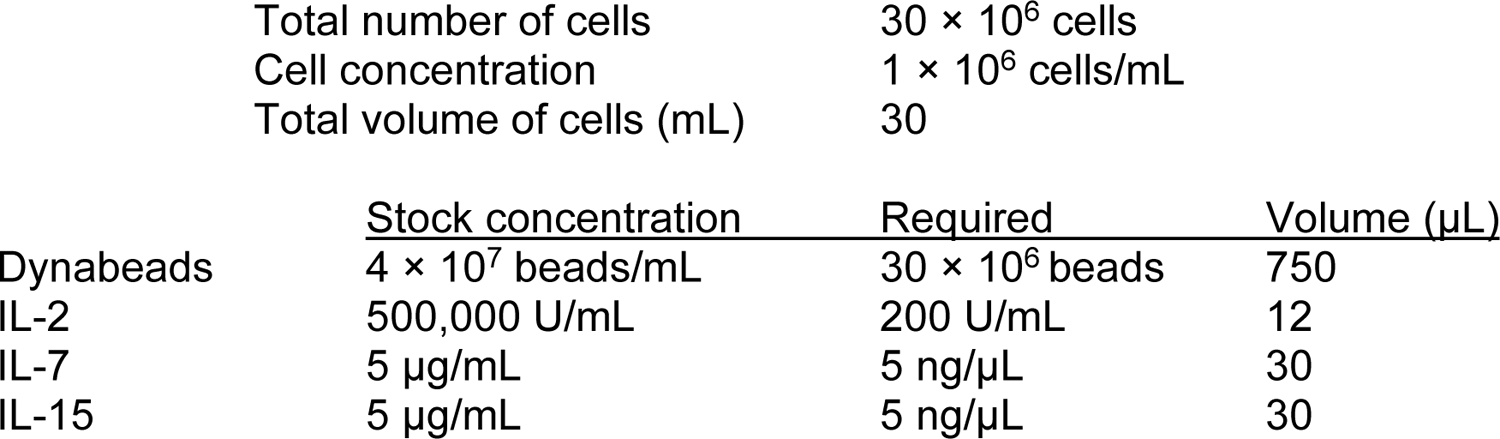
xxix. Vortex the Dynabead vial to mix the beads homogenously in solution. Pipette the required volume of beads from the stock bottle into a 15 mL tube and place the tube on the Dynamag^TM^-15. The beads will be drawn to the side of tube. Carefully pipette out the storage solution without disturbing the beads.
xxx. Remove the 15 mL tube from the magnetic stand and wash the beads with 1 mL DPBS. Using the same tip, pipette to resuspend the beads in DPBS.
xxxi. Repeat this wash step one more time, finally resuspending the beads in 1 mL growth medium set aside previously in Step xvii. **CRITICAL STEP** Dynabeads tend to settle at the bottom of the vial quickly, so make sure to vortex right before pipetting to ensure aliquoting from a homogenous suspension.
xxxii. Add the cytokines (IL-2, IL-7, and IL-15) into this 1 mL medium containing beads and mix well.
xxxiii. Take the 1 mL medium containing activation reagents and pipette into the 50 mL tube containing cells. Mix well and transfer to T25 flask.
xxxiv. Check cells under the microscope. Along with T cells in suspension, the smaller sized, brown magnetic beads should be present.
xxxv. Place cells back in the incubator set at 37°C and 5% CO2. *Day 0: Preparing cells for editing* **Timing 1.5 hours**
xxxvi. After 36–40 hours of activation, on the morning of the day of delivery, remove Dynabeads from cell culture.
xxxvii. Using a 10 mL disposable serological pipette, pipette cell suspension 10–20 times against the inner wall of the flask to break up the cell-bead clusters and make the cell suspension even.
xxxviii. Transfer cells into a 15 mL tube and place the tube on the magnet to separate the beads. In one swift motion, carefully pour out the cells into a clean new 50 mL tube, leaving behind the beads on the magnet. Repeat this step for the rest of the cell volume. **CRITICAL STEP** Be careful when pouring cells from the magnet into a tube and avoid spills.
xxxix. Take a 10 µL sample and count cells using a hemocytometer. Readjust concentration to 1 × 10^6^ cells/mL. **CRITICAL STEP ?Troubleshooting** Upon activation, T cells increase in size. This change is distinctly visible under the microscope when counting cells. Always make a note of how well the cells are activated on any given day. It is typical to attain 80–90% activation using Dynabeads. However, cells from some donors do not activate well, which may result in low editing efficiency.
xxxx. Centrifuge cells at 300 × *g* for 5 minutes RT, discard the supernatant, and resuspend cell pellet in fresh growth medium supplemented with 300 U/mL IL-2.
xxxxi. Aliquot 200 µL of prepared cell suspension (2 × 10^5^ cells) into each experimental well of a 96-well tissue culture treated plate. For T cells we recommend using U-bottom plates.
xxxxii. Return the cells to the incubator at 37°C and 5% CO2. **Pause point** Cells can be left in the incubator for approximately 2 days until PERC.

A. Culturing and preparing HSPCs for editing Timing 1 hour hands on, 2–3 days of culture

*Day −2: thaw frozen HSPCs* Timing 1 hour

i. Prepare HSPC culture medium according to instructions provided under ‘reagent setup’.
ii. Pre-warm 15 mL of SFEM II (for washing) and the required amount of HSPC culture medium in a 37°C water bath.
iii. Remove a cryovial of frozen cells from liquid nitrogen storage and transfer to dry ice.
iv. Wipe the outside of the vial with 70% ethanol and inside the biosafety cabinet, quarter turn the lid to release pressure and re-tighten.
v. Thaw the vial in a 37°C water bath. Gently swirl the vial and monitor thawing of the frozen cell stock, removing it from the water bath when 90% thawed.
vi. Wipe the vial again with 70% ethanol before placing it inside the BSC and transfer contents into a clean 50 mL conical tube.
vii. Wash the empty vial with 1 mL of pre-warmed non-supplemented SFEM II and slowly add this drop by drop to the 50 mL tube containing cells.
viii. Add the remaining non-supplemented SFEM II drop by drop into the 50 mL tube while gently swirling the cells (to slowly dilute out the DMSO in the freezing medium).
ix. Centrifuge at 300 × g for 10 minutes, RT to pellet the cells.
x. Aspirate the supernatant carefully, leaving behind ∼100 µL to not disturb the pellet. Flick the pellet to resuspend the cells.
xi. Resuspend the cells in HSPC culture medium to a cell concentration between 5 × 10^5^ – 1 × 10^6^ cells/mL for counting.
xii. Take a 10 µL sample and count cells using a hemocytometer. Readjust cell density to 2.5 × 10^5^ cells/mL with additional HSPC culture medium, and transfer cells into a labeled T25 or T75 flask as appropriate.
xiii. Lay the flask flat in an incubator at 37°C and 5% CO2. **Pause point** Cells should be cultured for 36–60 hours. **CRITICAL STEP** work quickly to maintain high cell viability. We do not recommend thawing more than 3–4 vials at the same time. *Day 0: Preparing cells for editing* Timing 1.5 hours
xiv. Pre-warm some SFEM II (for washing) and HSPC culture medium in a 37°C water bath.
xv. Resuspend the HSPCs and transfer the suspension to an appropriately sized tube. Wash the flask with SFEM II to collect any remaining cells and add to the tube.
xvi. Mix the cell suspension to evenly distribute the cells and take a 20 µL aliquot for counting. Count the number of cells and make a note of the viability. Dilute 20 µL cell suspension in 10 µL trypan blue to ensure the cells are concentrated enough to count at least 100 cells in four squares.
xvii. Centrifuge the cells at 300 × g for 10 minutes, RT to pellet the cells.
xviii. Aspirate the supernatant carefully so as to not disturb the pellet. Flick the pellet to resuspend the cells.
xix. Resuspend the cells in fresh, pre-warmed HSPC culture medium to 1.11 × 10^6^ cells/mL.
xx. Plate 90 µL of prepared cell suspension into each experimental well of a 96-well flat bottom TC-treated plate. Ensure the cell suspension remains well mixed during cell plating, to minimize cell number variation between wells. This step will result in 1 × 10^5^ cells per well.
xxi. Fill the bordering wells with 100–200 µL of 1× DPBS to minimize uneven evaporation of the experimental wells.
xxii. Return the cells to the incubator at 37°C and 5% CO2. **Pause point** Cells can be left in the incubator for several hours. We typically edit cells within 2–3 hours of plating, but have observed no issues with editing up to 7 hours post plating. ?Troubleshooting Assembling the CRISPR RNP Timing 1 hour
xxiii. Begin by cleaning the bench, tube rack, and pipettes with ELIMINase or RNase away, followed by 70% ethanol. **CRITICAL STEP** it is important to work with RNase-free solutions during this step, as RNase can hinder editing efficiency. We recommend using filter tips.
xxiv. Thaw gRNA on ice if frozen. Once thawed, place on the bench for 10 minutes to bring to room temperature.
xxv. Thaw Cas protein on ice. Once thawed, place on a tube rack for 10 minutes to bring to room temperature. **CRITICAL STEP** It is important to bring all reagents to room temperature before combining. This step can help prevent precipitation.
xxvi. We recommend making RNP at a maximum concentration of 12.5 µM, as it tends to precipitate when prepared at higher concentrations.
xxvii. Combine the required volume of Cas protein with RNP buffer. Mix by pipetting.
xxviii. In another 1.5 mL centrifuge tube, pipette the desired volume of gRNA.
xxix. Slowly mix the diluted Cas protein into the tube containing gRNA while stirring the pipette tip to mix. **CRITICAL STEP** Always add the Cas protein into a tube containing gRNA, and not the other way around. Adding gRNA into a tube containing Cas protein can result in precipitation.
xxx. Follow the table below to make 2.64 × 10^−9^ moles of RNP at a concentration of 12.5 µM. It is important to note that RNP amounts are based on the Cas protein component. To assemble 2.64 × 10^−9^ moles of RNP, one would need 2.64 × 10^−9^ moles of Cas protein and 3.96 × 10^−9^ moles of gRNA for a 1:1.5 molar ratio of protein:gRNA: **Pause point** Prepared RNP can be used to edit cells the same day and can be kept at room temperature for up to 4 hours. Alternatively, RNP can be flash-frozen and stored at −80°C for several weeks.

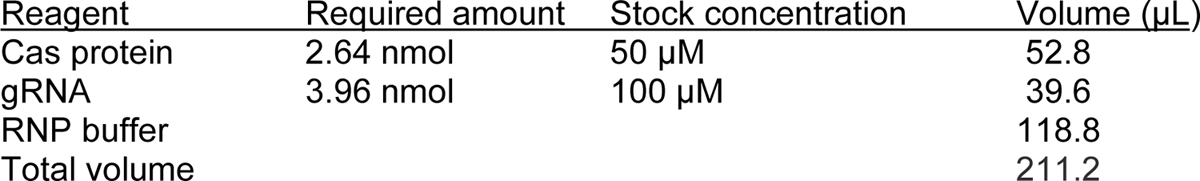 **Treatment of cells with editing reagents Timing 2 hours**
xxxi. The table below indicates example volumes of RNP, peptide, and buffer for an initial dose optimization experiment in T cells for a single well. We recommend scaling up the volumes to account for biological and/or technical replicates and pipetting loss.

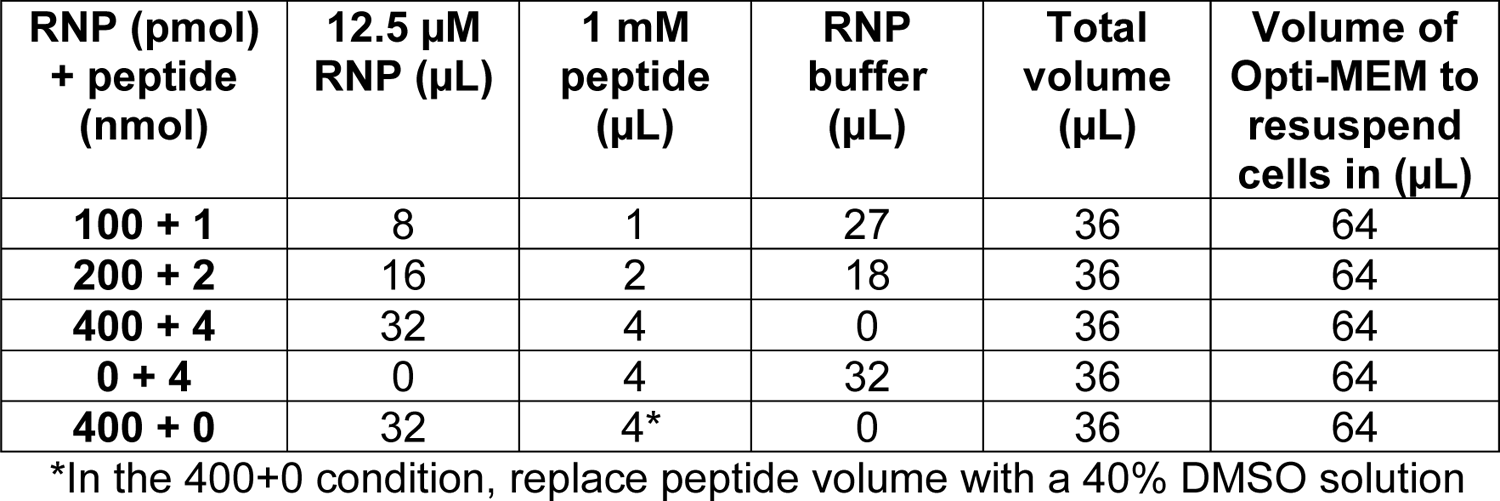

 A similar table can be followed for HSPCs, but we recommend limiting the volume of editing reagents to 10 µL to promote better viability. Concentrated RNP from Step 10 can be used for higher doses. To fit the PERC treatment in 10 µL, the peptide volume is fixed at 1 µL and the RNP volume is fixed at 9 µL. Different working stock concentrations of RNP and peptide should be prepared to test different PERC doses:

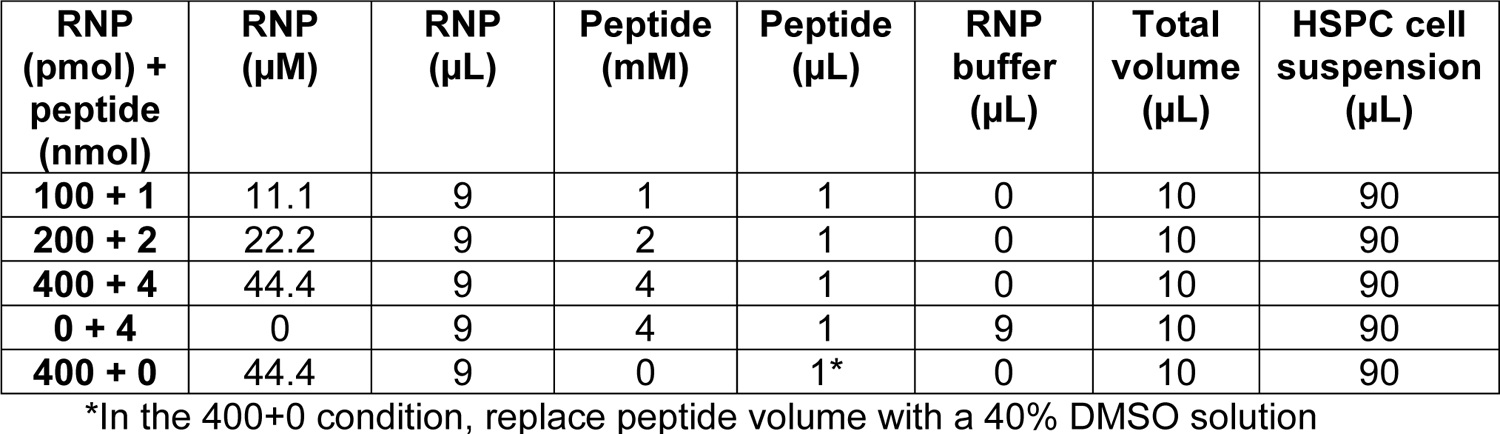
xxxii. If working with HSPCs, continue directly to step 14. For T cells, spin down the plate containing 2 × 10^5^ cells per well at 300 × g for 5 minutes at RT and aspirate the supernatant. If using a round-bottom plate, flick off the supernatant directly into a sink, lightly tapping the inverted plate once on a clean paper towel.
xxxiii. Resuspend the cells in an appropriate volume of Opti-MEM such that the total volume after adding editing reagents is 100 µL.
xxxiv. Make a 1–4 mM working solution of peptide in DEPC-treated water using the 10 mM DMSO stock prepared previously. For example, for a 1 mM working stock, mix 1 μL of 10 mM peptide and 9 μL of water, and pipette a few times until solution is homogenous. **CRITICAL STEP** The peptide working stock should be prepared fresh (on the same day).
xxxv. In the biosafety cabinet, mix peptide with RNP and incubate for 1–2 minutes. The solution may turn cloudy; pipette well until homogenous. **CRITICAL STEP** Do not allow RNP-peptide mixture to incubate for longer than 10 minutes. Minimize the incubation time as much as possible.
xxxvi. Bring the volume of all wells to the same total treatment volume by adding additional RNP buffer. This helps in easy multichannel pipetting of cells and editing reagents.
xxxvii. Mix the RNP-peptide formulation well via pipetting before aliquoting atop previously plated cells. **CRITICAL STEP** DO NOT fully pipette or mix PERC reagents and cells together. We have observed that pipetting of reagents and cells together lowers the editing efficiency. **?Troubleshooting**
xxxviii. Fill the bordering wells of the 96-well plate with 100 µL of sterile 1× DPBS to prevent uneven evaporation.
xxxix. Allow cells to incubate with the editing reagents in an incubator at 37°C for 1 hour. *For T cells*:
xxxx. During the above incubation, prepare a recovery medium as a 1.5x stock prepared according to the recipe described under reagent setup. Warm up the recovery medium in a water bath set at 37°C until needed.
xxxxi. After 1 hour of treatment, split the cell suspension in half by transferring ∼50 µL to a new plate. Using a multi-channel pipette, add 150 µL of 1.5× recovery medium on top of all the wells to bring the final volume to 200 µL. *For HSPCs*:
xxxxii. After 1 hour of treatment, split the cell suspension in half by transferring ∼50 µL to a new plate. Using a multi-channel pipette, add 150 µL of HSPC culture medium on top of all the wells to bring the final volume to 200 µL. *Optional:* To improve viability, we recommend washing off the PERC treatment after an hour and replacing with fresh supplemented medium
xxxxiii. Centrifuge cells at 300 × g for 5 minutes RT, discard the supernatant, and wash two times in 150 µL of warm non-supplemented medium, finally resuspending cells in their respective fresh prewarmed supplemented growth medium with cytokines. Fill empty wells with 1× DPBS.
xxxxiv. Return the cells to the incubator at 37°C and 5% CO2 **Pause point** 1-hour incubation of PERC reagents and cells. Editing cells for homology directed repair (HDR) **(Optional)** Timing 1 hour
xxxxv. After the 1-hour co-incubation of cells and PERC editing reagents, add 100 µL of non-supplemented medium on top, diluting the treatment 1:1.
xxxxvi. Transfer cells to a U- or V-bottom plate (if they are not already in one).
xxxxvii. Centrifuge the plate of cells at 300 × *g* for 5 minutes at RT, discard the supernatant, and resuspend cells in 100 µL of supplemented medium.
xxxxviii. For HSPCs, transfer cells back to a flat-bottom tissue culture treated 96-well plate.
xxxxix. If needed, dilute AAV in sterile 1× DPBS so that the desired MOI is in 5 µL per 100 µL of cells. Pipet the 5 µL directly into the treatment well. **CRITICAL STEP** DO NOT pipette or mix upon addition of AAV to cells. One might have to functionally titrate the AAV to determine an optimal MOI.
xxxxx. Place cells back in the incubator at 37°C and 5% CO2. Incubate the cells with AAV overnight. **Pause point** Incubate for 12–24 hours. Day 1: Wash off AAV:
xxxxxi. Add 100 µL of non-supplemented medium on top, diluting the treatment 1:1.
xxxxxii. Transfer cells to a U- or V-bottom plate if not already in one. Centrifuge cells at 300 × *g* for 5 minutes RT, discard the supernatant, and wash once more in 200 µL of warm non-supplemented medium.
xxxxxiii. Resuspend cells in 200 µL of supplemented medium and split the cells in half by transferring 100 µL into two wells each. Add 100 µL of warm supplemented medium on top of all the wells to bring the final volume to 200 µL. Day 1 or 2: Assessing cell viability by CellTiter-Glo assay (optional) Timing 1 hour
xxxxxiv. CellTiter-Glo is an easy high-throughput readout for measuring cell viability. Follow manufacturer’s instructions to assess viability of edited cells 1–2 days after delivery of editing reagents. We use as little as 20 μL of cell suspension with 20 μL of assembled kit reagent for this test.
xxxxxv. Alternatively, stain cells with trypan blue and assess viability by counting live and dead cells using a hemocytometer. Assessing editing efficiency
xxxxxvi. Editing efficiency can be assessed broadly at different levels. Follow option A for protein-level knockout assessment using flow cytometry and option B for gene-level editing assessed by sequencing.

See **Box 2** (next page) for recommendations on concentrating RNP if desired.

#### Box 2

##### Concentrating RNP (optional)

Whenever possible, we recommend that the editing reagent volume not exceed 10% of the media volume during co-incubation of cells and editing reagents. This aspect may be especially important for sensitive cells such as HSPCs. For example, in a 96-well plate with a total treatment volume of 100 µL per well, do not exceed 10 µL volume of editing reagents. To facilitate use of higher RNP doses without exceeding the recommended 10% treatment volume, one should prepare RNP at a higher concentration, which increases the risk of RNP aggregation and precipitation. We recommend initially preparing RNPs at a lower concentration and using centrifugal filter-based devices to concentrate RNP to the desired concentration. Below is an example for concentrating 11.1 µM up to 44.4 µM.

**CRITICAL STEP** Prepare an excess and expect a 20% loss of material.

1. For RNP concentration, use a filter with a 50 kDa MWCO (molecular weight cutoff), with 0.5 mL capacity.
2. Place a filter into a microcentrifuge tube (provided with the kit). Follow the manufacturer’s instructions for the correct positioning of the filter device.
3. Wash an Amicon Ultra centrifugal filter device with 400 µL RNP buffer.
4. **CRITICAL STEP** The pipette tip should not touch the membrane region of the Amicon Ultra centrifugal filter device. This can damage the filter, leading to loss of RNP due to leakage.
5. Spin at 13,000 × g for 5 minutes.
6. Invert the filter to collect the remaining liquid inside the filter in the microcentrifuge tube. Spin at 1,000 × g for 1 minute.
7. Transfer the prepared filter to a fresh microcentrifuge tube. Add 400 µL 11.1 µM RNP into the filter. There is a 400 µL mark on the Amicon filter. Set aside 5 µL of the input RNP for later.
8. Start with a 13,000 × *g* spin for 1 minute. Note the volume change following the spin.
9. **CRITICAL STEP** The spins should be performed in small increments to avoid overconcentrating the RNP.
10. Continue to spin until the volume reaches 100 µL on the Amicon filter.
11. **CRITICAL STEP** The RNP can become especially concentrated at the bottom of the filter. Redistribute the RNP by gently yet thoroughly pipetting it up and down between centrifugation runs to improve the flow.
12. **CRITICAL STEP** The rate of concentration will slow down as the volume decreases, so spin times will gradually increase.
13. Continue concentrating via 1–5 minute spins until the sample volume within the filter is slightly below the target volume. This should ensure the RNP is sufficiently concentrated.
14. Once the desired volume is reached, transfer and invert the filter containing the concentrated RNP into a fresh centrifuge tube.
15. Spin at 1,000 × g for 5 minutes to collect the concentrated RNP. Save the flow-through.
16. Transfer the concentrated RNP into a clean centrifuge tube.

Using a NanoDrop, compare the A280 values of the input RNP to the concentrated RNP. Adjust the concentrated RNP to the desired final concentration using the flow-through saved earlier. Expect to prepare about 80 μL concentrated RNP with a starting volume of 400 μL non-concentrated RNP.

### A. Day 4: Flow cytometry

#### Staining cells for flow cytometry Timing 1 hour

i. If using the recommended antibody panel below, prepare a staining solution by mixing antibodies at the given dilutions in flow cytometry buffer (see reagent preparation section for recipe). Scale volumes according to the experiment size. T cell staining solution:

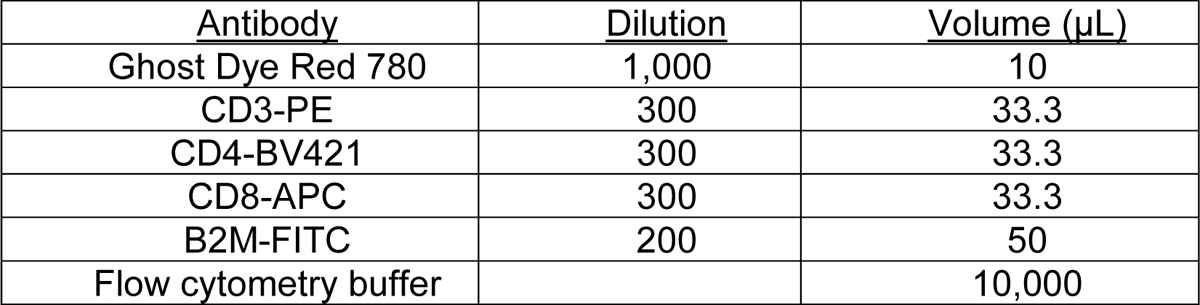 HSPC staining solution:

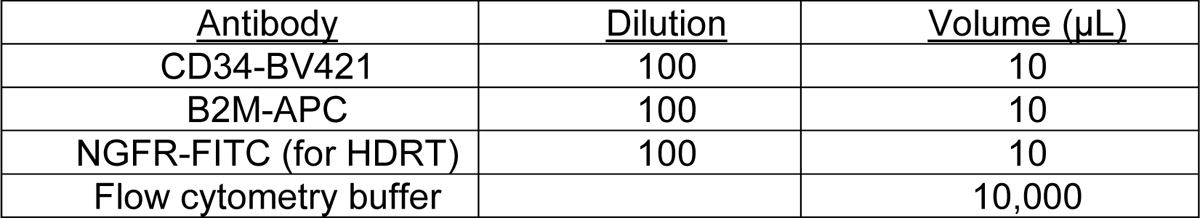 Using a viability dye for HSPCs is optional. We do not routinely use it because dead HSPCs appear in a different region of a flow plot of FSC vs. SSC.
ii. Centrifuge the plate containing cells at 300 × *g* for 5 min. If cells are in a flat-bottom plate, transfer to U- or V-bottom plates before centrifugation.
iii. Removing the lid of the 96-well plate, flick off media from the plate directly into a sink in one swift motion, and dab excess media onto a clean paper towel. Replace the lid of the plate immediately. These steps should be done quickly to avoid cell loss.
iv. Wash with 200 µL of flow cytometry buffer.
v. Repeat the previous two steps.
vi. Resuspend in 100 µL of staining solution prepared above in Step 34. Pipette to thoroughly mix the cells, and incubate at 4°C for 20 minutes in the dark.
vii. After incubation, centrifuge cells at 300 × g for 5 mins RT and flick off supernatant as described previously.
viii. Resuspend cells in 200 µL wash buffer.
ix. Centrifuge cells at 300 × g for 5 mins RT and flick off supernatant.
x. Resuspend cells in flow cytometry buffer, pipetting well to avoid cell clumping, and be gentle to avoid bubbles. The mixing step is best done right before running samples through the flow cytometer to produce a single-cell suspension.
xi. Run samples through a flow cytometer, following instrument-specific instructions. Export files into a folder of its own and if needed, and transfer to a computer with FlowJo software.
xii. Refer to **Fig. 7** for gating strategies for T cells or HSPCs. Export the plots and gated population cell counts in a table format into a spreadsheet and plot graphs to visualize the data. **?Troubleshooting**

**Figure 7:**
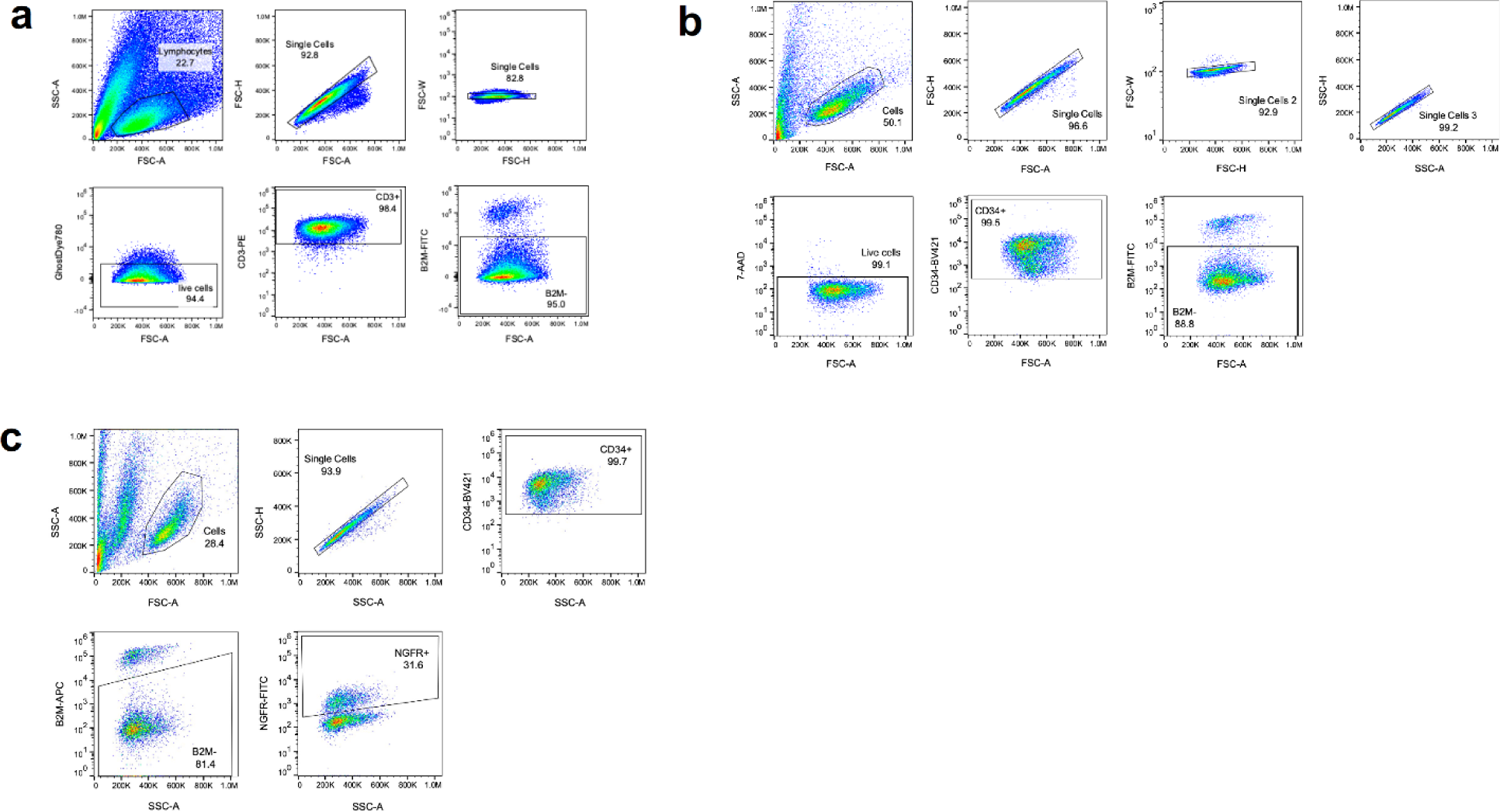
Gating strategies used for flow cytometry. **a.** Gating strategy for editing T cells using PERC. **b.** Gating strategy for editing HSPCs using PERC. **c.** Gating strategy for editing HSPCs using PERC when paired with HDRT.

### B. Day 3 or 4: Sequencing

#### Extracting genomic DNA for sequencing Timing 1 hour

i. Thaw Quick Extract (QE) DNA extraction solution for 15–20 minutes, to normalize it to RT. **CRITICAL STEP** QE solution should be stored at −20°C and aliquoted in smaller batches to prevent multiple freeze-thaws. An aliquot of QE solution may be stored at 4°C for up to a month.
ii. For HSPCs, resuspend the cells and transfer to a U-bottom or conical-bottom plate.
iii. Centrifuge the plate of cells at 300 × g for 5 minutes at RT, discard the supernatant, and wash with sterile DPBS.
iv. Pipette 20–50 µL of QE buffer directly on top of a single well of cell pellet and wait for 10 minutes.
v. Transfer cell lysates from 96-well plate to a PCR plate or tubes.
vi. Incubate the PCR plate in a thermocycler with the following conditions: 65°C for 10 minutes 95°C for 5 minutes 4°C hold
vii. Extracted genomic DNA from cells can be used as a template for PCR to amplify the region around the expected cut site of the gRNA to assess for indels. **Pause point** Genomic DNA can be used for PCR or stored at −20°C for later use. Preparing samples for sequencing Timing 4 hours
viii. Resuspend primers to 100 µM according to manufacturer’s recommendation. Stocks can be stored at −20°C for 1 year.
ix. Using the 100 µM stock, prepare a 10 µM working stock in nuclease-free water.
x. Thaw buffer, dNTPs, and primers (if frozen) on ice.
xi. Prepare a master mix using the Takara PrimeStar GXL polymerase kit according to the table below. Scale up the number of reactions according to the number of samples pertaining to the experiment. Include a negative control containing no DNA template (substitute with water), and account for 20% extra volume for pipetting loss. Primers used in our example experiment targeting *B2M* are provided in table 3 along with optimized annealing temperatures for Sanger sequencing or NGS.

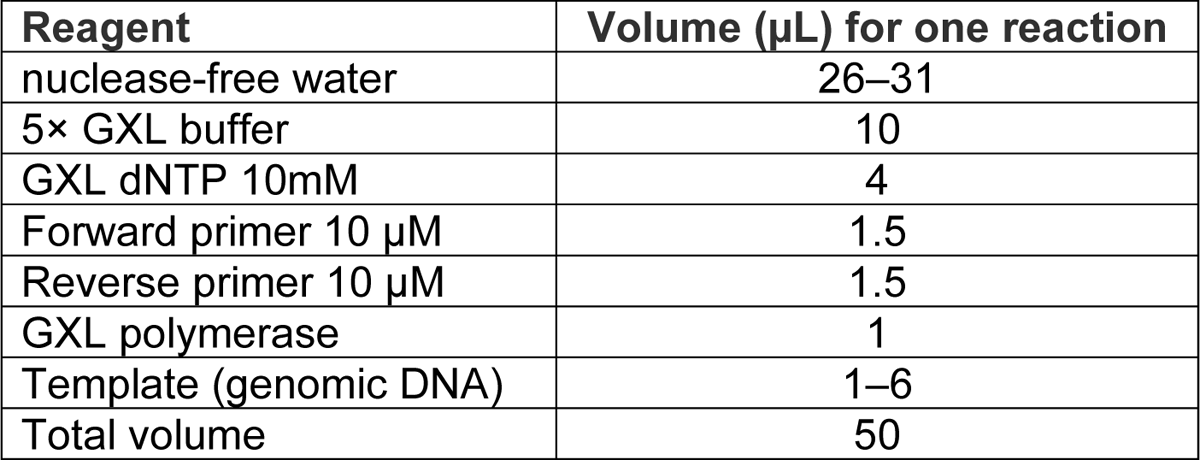 Thermocycler conditions:

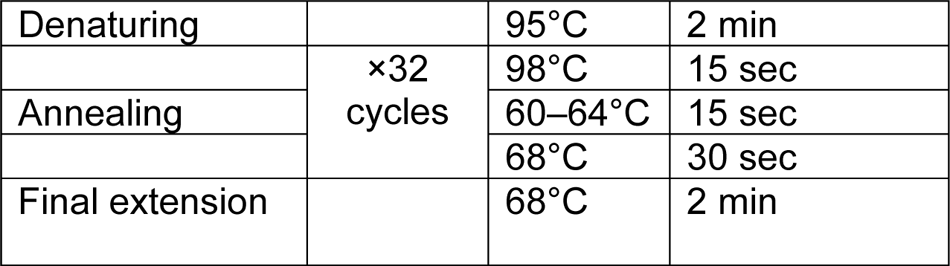
xii. Run 5 µL of PCR product on a 1% agarose gel to confirm the presence of the expected amplicon band in all of the samples. The negative control should not have an amplicon band. **Pause point** Samples can be stored at 4°C overnight or at −20°C for the long-term. **?Troubleshooting** **Magnetic bead-based clean-up of PCR products Timing 2 hours**
xiii. Add 1.8× the volume of SPRI beads to 1× PCR product (81 µL of beads to 45 µL of PCR product). Mix beads and PCR product thoroughly up to ten times by pipetting. Incubate for 10 minutes at RT.
xiv. Place PCR plate on a ring magnet for 10 minutes at RT. Amplicon-bound beads will form a ring around the magnet.
xv. Maintaining the plate on the magnet, aspirate the supernatant and wash the beads with 180 µL of freshly made 70% ethanol. **CRITICAL STEP** Only use freshly mixed up 70% ethanol. Avoid scraping the beads with the pipette tips when washing.
xvi. Wait 30 seconds before aspirating the ethanol. Repeat the wash for a total of two times.
xvii. Air-dry for 2–5 minutes. Remove any ethanol at the bottom of the well by aspirating using a 20 µL filter tip. Beads should change appearance from glossy/wet to matte/dry. **CRITICAL STEP** If the wait time is too long, the beads will start to dry and will be difficult to resuspend. But at the same time, it is important to ensure little to no ethanol is left behind before elution.
xviii. Elute DNA in at least 30 µL nuclease-free water, mix well, and incubate for at least 15 minutes at RT. **Pause point** Samples can be left to elute for longer, up to one hour on the bench or several hours to overnight at 4°C.
xix. Place PCR plate on the magnet to separate beads and recover eluted clean PCR amplicon in the supernatant by transferring 20 µL of to a fresh clean PCR plate. **CRITICAL STEP** Ensure beads do not get transferred over along with supernatant, which can happen if trying to transfer the entire volume.
xx. Quantify amplicon on a Nanodrop and confirm purity by assessing the A260/A280 ratio and/or optionally by running 1 µL of sample on a 1% agarose gel. **?Troubleshooting** Next-generation sequencing (optional)
xxi. If you have access to a sequencing core or an Illumina sequencer, proceed according to their instructions for amplicon sequencing. We used Illumina MiSeq 300bp paired-end reads and recommend a depth of at least 10,000 reads.
xxii. Visit the CRISPResso2^49^ website (http://crispresso2.pinellolab.org/submission) and follow the instructions to upload the required files and information to analyze data. Results can be downloaded in HTML or PDF format.
xxiii. For analyzing multiple samples at once for the same amplicon, CRISPResso2 can be used. But for more than one target amplicon, we recommend using Cortado (https://github.com/staciawyman/cortado). Sanger sequencing (optional)
xxiv. Submit purified PCR amplicons for Sanger sequencing according to the vendor’s instructions.
xxv. Download sequencing files provided. **?Troubleshooting** Analysis of sequencing data ICE analysis
xxvi. Visit https://ice.synthego.com and choose the ‘sample by sample upload’ option for analysis of a small set of samples. Fill in the details requested on the portal by providing a unique name to the sample, the gRNA sequence without the PAM, and the HDRT sequence for knock-in experiments.
xxvii. For ‘control file’, upload the sequence file corresponding to a non-treated or non-edited sample, and upload the edited sample sequence file to the ‘experiment file’ section. Click on ‘add sample to analysis’.
xxviii. Repeat steps 73–74 to add another sample to the analysis and/or proceed by clicking on the ‘analyze’ button.
xxix. If there are a large number of samples, we recommend using the ‘batch upload’ method. Download the template/example files from the ICE analysis home page and use their ‘template_ice_definitions’ file as a template to provide details for analysis.
xxx. Collect all .ab1 sequence files into one folder, making sure file names (including file format) exactly match the sample names provided on the template file. Compress the folder to a .zip file and upload to the ICE portal along with the template excel sheet. Click ‘analyze’ and wait up to several minutes for analysis to be completed.
xxxi. Data are available for visualization on the web page and can be downloaded as an excel sheet. Use the “ICE d” score readout for the closest matching value to indel-level readout using NGS. **?Troubleshooting** TIDE analysis
xxxii. Plot a graph using the ICE editing efficiencies to visualize the editing results.
xxxiii. Alternatively, Tracking of INDELs by decomposition (TIDE) analysis can be used. Visit https://tide.nki.nl/ and follow instructions provided on the website for analysis^70^.
xxxiv. Paste the gRNA sequence (without the PAM sequence) in the box provided.
xxxv. Upload the control and test samples. If there is more than one test sample, use the batch version and upload multiple .ab1 or .scn files.
xxxvi. Click on the results or decomposition tab on the right, click on ‘generate/refresh downloads’. A progress bar at the bottom of the screen will show processing in progress. Once finished, click on the ‘download plots’ button to download either a .pdf or .csv file with the results.
xxxvii. For quantification of knock-in, use the TIDER tool.

#### TIMELINE

Step 1A culturing and preparing primary human T cells: 3 h hands-on, 3 d of culture

Step 1B culturing and preparing HSPCs: 1h hands-on, 2 d of culture

Step 2–9 assembly of CRISPR RNP: 1 h

Step 10–23 treatment of cells with editing reagents: 2 h, 1 h incubation, 3–4 d in culture

Step 24–32 AAV mediated HDR: 1 h hands-on, 1 h incubation, 1 d in culture

Step 33–34 assessing cell viability: 1 h

Step 35A Assessing editing efficiency by flow cytometry: 1–2 h hands-on

Step 35B Assessing editing efficiency by sequencing

Step 35B i-vii Extracting genomic DNA for sequencing: 1 h

Step 35B viii-xx-preparing samples for sequencing: 2 h hands-on

Step 35B xxi-xxiii Next generation sequencing: 3–7 d turnaround

Step 35B xxiv-xxv Sanger sequencing: 1 d turnaround

Step 35B xxvi-xxxvii Analysis of sequencing data: 1–2 h hands-on

## Author contributions

S.U.S., M.C., J.J.M., L.D., D.N.N., J.E., and R.C.W. conceived the study. S.U.S. and M.C. designed the experiments, performed experiments, and analyzed data. J.J.M. and K.A. conceived and prepared reagents. L.D. designed the experiments. S.K.W. analyzed data. N.K. performed experiments. S.U.S., M.C., J.J.M., D.N.N., J.E. and R.C.W. wrote and edited the manuscript. All authors read the manuscript and agree to its contents.

## Acknowledgments

We thank C. Jeans of the QB3 Berkeley MacroLab for assistance. Sarah Pyle and the study authors adapted CRISPR enzyme illustrations from the Innovative Genomics Institute Glossary Icon Collection, originally by Christine Liu of Two Photon Art. S.U.S. and R.C.W. are supported by NIH award UG3AI150552 as part of the Somatic Cell Genome Editing (SCGE) consortium. D.N.N. is supported by NIH grants L40AI140341 and K08AI153767, and the UCSF LGR Innovation Award.

## Data availability

Data supporting this manuscript are available from the corresponding author upon reasonable request. Cas12a-Ultra-5xNLS plasmid is available on Addgene (#218775). Sequencing data will be made available from the NCBI SRA.

## REFERENCES

1. Taha, E. A., Lee, J. & Hotta, A. Delivery of CRISPR-Cas tools for *in vivo* genome editing therapy: Trends and challenges. J. Controlled Release 342, 345–361 (2022).

2. Sheridan, C. The world’s first CRISPR therapy is approved: who will receive it? Nat. Biotechnol. 42, 3–4 (2023).

3. Stadtmauer, E. A. et al. CRISPR-engineered T cells in patients with refractory cancer. Science 367, eaba7365 (2020).

4. Chiesa, R. et al. Base-Edited CAR7 T Cells for Relapsed T-Cell Acute Lymphoblastic Leukemia. N. Engl. J. Med. 389, 899–910 (2023).

5. Bhokisham, N. et al. CRISPR-Cas System: The Current and Emerging Translational Landscape. Cells 12, 1103 (2023).

6. Liu, X. et al. Advances in CRISPR/Cas gene therapy for inborn errors of immunity. Front. Immunol. 14, 1111777 (2023).

7. Zhang, S. et al. Current trends of clinical trials involving CRISPR/Cas systems. Front. Med. 10, 1292452 (2023).

8. Shifrut, E. et al. Genome-wide CRISPR Screens in Primary Human T Cells Reveal Key Regulators of Immune Function. Cell 175, 1958–1971.e15 (2018).

9. Schmidt, R. et al. Base-editing mutagenesis maps alleles to tune human T cell functions. Nature 625, 805–812 (2024).

10. Tothova, Z. et al. Multiplex CRISPR/Cas9-Based Genome Editing in Human Hematopoietic Stem Cells Models Clonal Hematopoiesis and Myeloid Neoplasia. Cell Stem Cell 21, 547–555.e8 (2017).

11. Canver, M. C. et al. BCL11A enhancer dissection by Cas9-mediated in situ saturating mutagenesis. Nature 527, 192–197 (2015).

12. Ravi, N. S. et al. Identification of novel HPFH-like mutations by CRISPR base editing that elevate the expression of fetal hemoglobin. eLife 11, e65421 (2022).

13. Cromer, M. K. et al. Ultra-deep sequencing validates safety of CRISPR/Cas9 genome editing in human hematopoietic stem and progenitor cells. Nat. Commun. 13, 4724 (2022).

14. Del’Guidice, T. et al. Membrane permeabilizing amphiphilic peptide delivers recombinant transcription factor and CRISPR-Cas9/Cpf1 ribonucleoproteins in hard-to-modify cells. PLOS ONE 13, e0195558 (2018).

15. Foss, D. V. et al. Peptide-mediated delivery of CRISPR enzymes for the efficient editing of primary human lymphocytes. Nat. Biomed. Eng. 7, 647–660 (2023).

16. Zhang, Z. et al. Efficient engineering of human and mouse primary cells using peptide-assisted genome editing. Nat. Biotechnol. 42, 305–315 (2023).

17. Rouet, R. et al. Receptor-Mediated Delivery of CRISPR-Cas9 Endonuclease for Cell-Type-Specific Gene Editing. J. Am. Chem. Soc. 140, 6596–6603 (2018).

18. Krishnamurthy, S. et al. Engineered amphiphilic peptides enable delivery of proteins and CRISPR-associated nucleases to airway epithelia. Nat. Commun. 10, 4906 (2019).

19. Kulhankova, K., et al. Shuttle peptide delivers base editor RNPs to rhesus monkey airway epithelial cells in vivo. Nat. Commun. 14, 8051 (2023).

20. Algayer, B. et al. Novel pH Selective, Highly Lytic Peptides Based on a Chimeric Influenza Hemagglutinin Peptide/Cell Penetrating Peptide Motif. Molecules 24, 2079 (2019).

21. Bak, R. O., Dever, D. P. & Porteus, M. H. CRISPR/Cas9 genome editing in human hematopoietic stem cells. Nat. Protoc. 13, 358–376 (2018).

22. Romero, Z. et al. Editing the Sickle Cell Disease Mutation in Human Hematopoietic Stem Cells: Comparison of Endonucleases and Homologous Donor Templates. Mol. Ther. J. Am. Soc. Gene Ther. 27, 1389–1406 (2019).

23. Nguyen, D. N. et al. Polymer-stabilized Cas9 nanoparticles and modified repair templates increase genome editing efficiency. Nat. Biotechnol. 38, 44–49 (2020).

24. Vavassori, V. et al. Lipid nanoparticles allow efficient and harmless ex vivo gene editing of human hematopoietic cells. Blood 142, 812–826 (2023).

25. Bothmer, A. et al. Detection and Modulation of DNA Translocations During Multi-Gene Genome Editing in T Cells. CRISPR J. 3, 177–187 (2020).

26. Kim, S., Kim, D., Cho, S. W., Kim, J. & Kim, J.-S. Highly efficient RNA-guided genome editing in human cells via delivery of purified Cas9 ribonucleoproteins. Genome Res. 24, 1012–1019 (2014).

27. Sheridan, C. Off-the-shelf, gene-edited CAR-T cells forge ahead, despite safety scare. Nat. Biotechnol. 40, 5–8 (2022).

28. Bashor, C. J., Hilton, I. B., Bandukwala, H., Smith, D. M. & Veiseh, O. Engineering the next generation of cell-based therapeutics. Nat. Rev. Drug Discov. 21, 655–675 (2022).

29. Ardeniz, Ö. et al. β2-Microglobulin deficiency causes a complex immunodeficiency of the innate and adaptive immune system. J. Allergy Clin. Immunol. 136, 392–401 (2015).

30. Eyquem, J. et al. Targeting a CAR to the TRAC locus with CRISPR/Cas9 enhances tumour rejection. Nature 543, 113–117 (2017).

31. Rezalotfi, A., Fritz, L., Förster, R. & Bošnjak, B. Challenges of CRISPR-Based Gene Editing in Primary T Cells. Int. J. Mol. Sci. 23, 1689 (2022).

32. Schumann, K. et al. Generation of knock-in primary human T cells using Cas9 ribonucleoproteins. Proc. Natl. Acad. Sci. U. S. A. 112, 10437–10442 (2015).

33. DiTommaso, T. et al. Cell engineering with microfluidic squeezing preserves functionality of primary immune cells in vivo. Proc. Natl. Acad. Sci. U. S. A. 115, E10907–E10914 (2018).

34. Philippidis, A. CASGEVY Makes History as FDA Approves First CRISPR/Cas9 Genome Edited Therapy. Hum. Gene Ther. 35, 1–4 (2024).

35. Schmiderer, L. et al. Efficient and nontoxic biomolecule delivery to primary human hematopoietic stem cells using nanostraws. Proc. Natl. Acad. Sci. U. S. A. 117, 21267–21273 (2020).

36. Martínez Bedoya, D., Dutoit, V. & Migliorini, D. Allogeneic CAR T Cells: An Alternative to Overcome Challenges of CAR T Cell Therapy in Glioblastoma. Front. Immunol. 12, 640082 (2021).

37. Germino-Watnick, P. et al. Hematopoietic Stem Cell Gene-Addition/Editing Therapy in Sickle Cell Disease. Cells 11, 1843 (2022).

38. Mock, U. et al. Automated manufacturing of chimeric antigen receptor T cells for adoptive immunotherapy using CliniMACS prodigy. Cytotherapy 18, 1002–1011 (2016).

39. Gotti, E. et al. Optimization of therapeutic T cell expansion in G-Rex device and applicability to large-scale production for clinical use. Cytotherapy 24, 334–343 (2022).

40. Gillmore, J. D. et al. CRISPR-Cas9 In Vivo Gene Editing for Transthyretin Amyloidosis. N. Engl. J. Med. 385, 493–502 (2021).

41. Wei, T., Cheng, Q., Min, Y.-L., Olson, E. N. & Siegwart, D. J. Systemic nanoparticle delivery of CRISPR-Cas9 ribonucleoproteins for effective tissue specific genome editing. Nat. Commun. 11, 3232 (2020).

42. Cheng, Q. et al. Selective organ targeting (SORT) nanoparticles for tissue-specific mRNA delivery and CRISPR-Cas gene editing. Nat. Nanotechnol. 15, 313–320 (2020).

43. Miao, L., Zhang, Y. & Huang, L. mRNA vaccine for cancer immunotherapy. Mol. Cancer 20, 41 (2021).

44. Baden, L. R. et al. Efficacy and Safety of the mRNA-1273 SARS-CoV-2 Vaccine. N. Engl. J. Med. 384, 403–416 (2021).

45. Wang, X. et al. Preparation of selective organ-targeting (SORT) lipid nanoparticles (LNPs) using multiple technical methods for tissue-specific mRNA delivery. Nat. Protoc. 18, 265–291 (2023).

46. Maeki, M., Uno, S., Niwa, A., Okada, Y. & Tokeshi, M. Microfluidic technologies and devices for lipid nanoparticle-based RNA delivery. J. Control. Release Off. J. Control. Release Soc. 344, 80–96 (2022).

47. Lin, S.-W., Nguyen, V. Q. & Lin, S. Preparation of Cas9 Ribonucleoproteins for Genome Editing. Bio-Protoc. 12, e4420 (2022).

48. Coin, I., Beyermann, M. & Bienert, M. Solid-phase peptide synthesis: from standard procedures to the synthesis of difficult sequences. Nat. Protoc. 2, 3247–3256 (2007).

49. Clement, K. et al. CRISPResso2 provides accurate and rapid genome editing sequence analysis. Nat. Biotechnol. 37, 224–226 (2019).

50. Ran, F. A. et al. Genome engineering using the CRISPR-Cas9 system. Nat. Protoc. 8, 2281–2308 (2013).

51. Hendel, A. et al. Chemically modified guide RNAs enhance CRISPR-Cas genome editing in human primary cells. Nat. Biotechnol. 33, 985–989 (2015).

52. Rathbone, T. et al. Electroporation-Mediated Delivery of Cas9 Ribonucleoproteins Results in High Levels of Gene Editing in Primary Hepatocytes. CRISPR J. 5, 397–409 (2022).

53. Mir, A. et al. Heavily and fully modified RNAs guide efficient SpyCas9-mediated genome editing. Nat. Commun. 9, 2641 (2018).

54. Kim, S. et al. CRISPR RNAs trigger innate immune responses in human cells. Genome Res. 28, 367–373 (2018).

55. Zhang, T., Gao, Y., Wang, R. & Zhao, Y. Production of Guide RNAs in vitro and in vivo for CRISPR Using Ribozymes and RNA Polymerase II Promoters. Bio-Protoc. 7, e2148 (2017).

56. Wu, Y. et al. Highly efficient therapeutic gene editing of human hematopoietic stem cells. Nat. Med. 25, 776–783 (2019).

57. Métais, J.-Y. et al. Genome editing of HBG1 and HBG2 to induce fetal hemoglobin. Blood Adv. 3, 3379–3392 (2019).

58. Stahl, E. C. et al. Genome editing in the mouse brain with minimally immunogenic Cas9 RNPs. Mol. Ther. J. Am. Soc. Gene Ther. 31, 2422–2438 (2023).

59. Staahl, B. T. et al. Efficient genome editing in the mouse brain by local delivery of engineered Cas9 ribonucleoprotein complexes. Nat. Biotechnol. 35, 431–434 (2017).

60. Zhang, L. et al. AsCas12a ultra nuclease facilitates the rapid generation of therapeutic cell medicines. Nat. Commun. 12, 3908 (2021).

61. Luk, K. et al. Optimization of Nuclear Localization Signal Composition Improves CRISPR-Cas12a Editing Rates in Human Primary Cells. GEN Biotechnol. 1, 271–284 (2022).

62. Kleinstiver, B. P. et al. Engineered CRISPR-Cas12a variants with increased activities and improved targeting ranges for gene, epigenetic and base editing. Nat. Biotechnol. 37, 276–282 (2019).

63. Fang, D., Wang, R., Yu, X. & Tian, Y. Construction of Cyclic Cell-Penetrating Peptides for Enhanced Penetration of Biological Barriers. J. Vis. Exp. JoVE (2022) doi:10.3791/64293.

64. Fripont, S., Marneffe, C., Marino, M., Rincon, M. Y. & Holt, M. G. Production, Purification, and Quality Control for Adeno-associated Virus-based Vectors. JoVE J. Vis. Exp. e58960 (2019) doi:10.3791/58960.

65. McClure, C., Cole, K. L. H., Wulff, P., Klugmann, M. & Murray, A. J. Production and Titering of Recombinant Adeno-associated Viral Vectors. JoVE J. Vis. Exp. e3348 (2011) doi:10.3791/3348.

66. Grieger, J. C., Choi, V. W. & Samulski, R. J. Production and characterization of adeno-associated viral vectors. Nat. Protoc. 1, 1412–1428 (2006).

67. Shy, B. R. et al. High-yield genome engineering in primary cells using a hybrid ssDNA repair template and small-molecule cocktails. Nat. Biotechnol. 41, 521–531 (2023).

68. Okamoto, S., Amaishi, Y., Maki, I., Enoki, T. & Mineno, J. Highly efficient genome editing for single-base substitutions using optimized ssODNs with Cas9-RNPs. Sci. Rep. 9, 4811 (2019).

69. Conant, D. et al. Inference of CRISPR Edits from Sanger Trace Data. CRISPR J. 5, 123–130 (2022).

70. Brinkman, E. K., Chen, T., Amendola, M. & van Steensel, B. Easy quantitative assessment of genome editing by sequence trace decomposition. Nucleic Acids Res. 42, e168 (2014).

71. Brinkman, E. K. et al. Easy quantification of template-directed CRISPR/Cas9 editing. Nucleic Acids Res. 46, e58 (2018).

72. Kluesner, M. G. et al. EditR: A Method to Quantify Base Editing from Sanger Sequencing. CRISPR J. 1, 239–250 (2018).

73. Tsuchida, C. A. et al. Mitigation of chromosome loss in clinical CRISPR-Cas9-engineered T cells. BioRxiv Prepr. Serv. Biol. 2023.03.22.533709 (2023) doi:10.1101/2023.03.22.533709.

74. Zeng, J. et al. Gene editing without ex vivo culture evades genotoxicity in human hematopoietic stem cells. bioRxiv 2023.05.27.542323 (2023) doi:10.1101/2023.05.27.542323.

75. Philippidis, A. Graphite Bio Pauses Lead Gene Editing Program in Sickle Cell Disease. Hum. Gene Ther. 34, 90–93 (2023).

76. Oh, S. A. et al. High-efficiency nonviral CRISPR/Cas9-mediated gene editing of human T cells using plasmid donor DNA. J. Exp. Med. 219, e20211530 (2022).

77. Hoerster, K. et al. HLA Class I Knockout Converts Allogeneic Primary NK Cells Into Suitable Effectors for ‘Off-the-Shelf’ Immunotherapy. Front. Immunol. 11, 586168 (2020).

78. Koressaar, T. & Remm, M. Enhancements and modifications of primer design program Primer3. Bioinforma. Oxf. Engl. 23, 1289–1291 (2007).

79. Untergasser, A. et al. Primer3--new capabilities and interfaces. Nucleic Acids Res. 40, e115 (2012).

80. Kõressaar, T. et al. Primer3_masker: integrating masking of template sequence with primer design software. Bioinforma. Oxf. Engl. 34, 1937–1938 (2018).

81. Ye, J. et al. Primer-BLAST: a tool to design target-specific primers for polymerase chain reaction. BMC Bioinformatics 13, 134 (2012).

82. Protocol for Isolating Mononuclear Cells from Whole Blood. https://www.stemcell.com/isolating-mononuclear-cells-from-whole-blood-by-density-gradient-centrifugation.html.

